# Pathogenic human autoantibodies target the soluble CASPR2 ectodomain to induce network hyperexcitability

**DOI:** 10.64898/2026.11.16.766757

**Authors:** Débora Serrenho, Nuno del Villar, Vitor Bueno, Dominique Fernandes, Quin Tutor-New, Takahisa Kanekiyo, Elinor Wing, Robyn Williams, Stephan Steinke, Luís F. Ribeiro, Paulo Pinheiro, Ester Coutinho, Sarosh R Irani, Ana Luísa Carvalho

## Abstract

The mechanisms by which autoantibodies contribute to antibody-mediated neurological diseases remain poorly understood. Autoantibodies targeting the extracellular domain of CASPR2 are found in patients with clinical features consistent with hyperexcitability, including neuromyotonia and seizures. These antibodies are reported to disrupt the functions of membrane-embedded CASPR2. However, given that CASPR2 is also detected in cerebrospinal fluid (CSF), which contains CASPR2 autoantibodies, targeting of CSF-shed CASPR2 may represent an additional pathogenic mechanism, introducing a novel paradigm of soluble antigen modulation by autoantibodies. Here, we found that patient-derived polyclonal and monoclonal CASPR2 antibodies enhanced neuronal network excitability and selectively amplified excitatory synaptic transmission in human and rat primary cortical neurons. In addition, CASPR2 antibodies impaired calcium homeostasis and altered the intrinsic properties of both excitatory and inhibitory neurons. Notably, these antibodies depleted CASPR2 from both neuronal culture medium and patient CSF, suggesting targeting of the shed CASPR2 ectodomain. Knockdown of plasma membrane calcium ATPase 2 (PMCA2), a reported target of the CASPR2 ectodomain, abrogated the observed effects of CASPR2 antibodies. Together, our findings identify shed CASPR2 as a underrecognized pathogenic target of CASPR2 autoantibodies and demonstrate that its neutralization disrupts PMCA2-mediated calcium regulation, driving neuronal network hyperexcitability. More broadly, we identify the neuronal sheddome as a source of autoantibody targets and soluble protein neutralization as a novel disease mechanism, highlighting shed CASPR2 as a potential biomarker and therapeutic target.

**One Sentence Summary:** The soluble CASPR2 ectodomain is targeted by human antibodies, revealing soluble-protein interference as a pathogenic mechanism in CASPR2-associated diseases.

## INTRODUCTION

Autoantibodies targeting extracellular epitopes of neuroglial antigens are typically considered pathogenic because these epitopes are directly accessible to circulating and cerebrospinal fluid (CSF) antibodies (*1, 2*). To date, mechanistic research has centered on autoantibody-induced dysfunction of the antigenic target via mechanisms including cross-linking, internalization, functional interference or by disruption of interactions with key partners (*2*). Less attention has been given to the downstream cellular and molecular mechanisms that link target disruption to the resulting clinical symptoms. Yet, such downstream mechanisms could uncover novel therapeutic targets, a major unmet need given that many patients continue to experience substantial residual deficits (*1, 3–7*).

Antibodies against Contactin-Associated Protein-like 2 (CASPR2) are detected in the serum and CSF of patients presenting a wide spectrum of neurological symptoms involving both the peripheral nervous system (PNS) and central nervous system (CNS). These include peripheral nerve hyperexcitability, neuropathic pain, cognitive dysfunction, seizures and movement disorders (*4, 8–11*). Passive transfer models using patient-derived polyclonal and monoclonal antibodies have established the pathogenicity of CASPR2-antibodies, with these antibodies inducing cognitive dysfunction and pain (*12–15*). Recognised molecular mechanisms have included internalisation of CASPR2 (*15, 16*) and modulation of crosstalk with interacting proteins such as Contactin 2, Kv channels (*13, 14*) and AMPA receptors (*13, 15, 17*). Yet, the dominant molecular mechanisms and the downstream cellular mechanisms underlying the patient’s clinical manifestations remained elusive. This constitutes a barrier to better understand disease pathogenesis, improve therapeutic strategy and long-term outcomes. Importantly, although patients often have favourable initial responses to immunotherapies, they frequently relapse and can develop medication side effects or long-term sequelae (*1, 2, 4, 5, 11, 18–20*).

CASPR2 is a cell adhesion molecule with a single transmembrane domain, and a large extracellular domain (ectodomain) that represents the target of patient antibodies. CASPR2 is expressed in both axons and synapses of excitatory and inhibitory neurons, where it contributes to the regulation of neuronal excitability and synaptic function (*14, 14, 17, 21–26*). However, recent studies have demonstrated that the CASPR2 ectodomain undergoes proteolytic shedding and is released into the CSF (*27, 28*). This shed ectodomain is reported to exert physiological effects by binding to the plasma membrane calcium ATPase PMCA2 to enhance calcium extrusion and reduce neuronal network activity. These observations suggest that the shed CASPR2 ectodomain functions as an endogenous regulator of neuronal excitability. They also raise the possibility that pathogenic antibodies may target not only membrane-bound CASPR2 but also its physiologically active soluble ectodomain, and contribute to disease pathogenesis.

Here, we tested the hypothesis that patient-derived CASPR2-antibodies exert pathogenic effects via the soluble CASPR2 ectodomain and interfere with its physiological function. Using primary rat cortical neurons and patient CSF, we demonstrate that patient-derived CASPR2-antibodies bind the soluble CASPR2 ectodomain, depleting free soluble CASPR2 which triggers PMCA2-mediated alterations in calcium dynamics that induce neuronal network hyperexcitability. Our findings identify a previously unrecognised mechanism of human autoantibodies, offering opportunities for precision targeting therapeutics.

## RESULTS

### CASPR2 antibodies induce increased network excitability in primary rat cultured neurons

To test how CASPR2 Abs might affect neuronal excitability, we used IgGs purified (pIgGs) from the plasma of three control individuals and three patients with high-titre CASPR2 Abs. Clinical features of the three patients, their antibody titers and isotype subclasses are summarized in Table S1. Antibody specificity and binding of patient, but not healthy control, sera was confirmed using a tissue-based assay (TBA, dilution 1:100, Fig. S1A) and live cell-based assays in cultured neurons and CASPR2-transfected HEK cells (LCBA, dilution 1:100, Fig. S1B,C). All patient failed to bind HEK transfected with other antigens, including LGI1 and AQP4. Hence, patient IgGs specifically bound the extracellular domain of CASPR2.

Since seizures arise from excessive hypersynchronous neuronal network activity, we first asked whether these antibodies alter spontaneous neuronal network activity. We used Microelectrode Array (MEA) recordings in both hiPSC derived excitatory neurons and primary neurons of the rat cortex, a brain region with high CASPR2 expression (*29, 30*). Mature neuronal cultures were treated with sera (1:100 dilutions) or individual pIgGs (Fig. 1A,B) at 20 µg/mL, both to mimic physiological IgG levels in the brain (*31, 32*). In rat neurons, by comparison to healthy controls, patient pIgGs induced network hyperexcitability after 5 days, as evidenced by increases in Area Under Cross-Correlation, Weighted Mean Firing Rate, Burst Frequency, and Network Burst Frequency (Fig. 1C,D). Analysis of network activity over time revealed that Patient pIgGs increased network firing within the first hours of incubation, with the effect progressively increasing over time (Fig. 1E-G). Notably, these changes were consistent across individual patient samples (Fig. S2A-C). Additionally, in human iNeurons (Fig. S3), after 6 h incubations with two CASPR2-antibody encephalitis patient sera, we observed increased firing rates by comparison to sera from a healthy control. Taken together, these results indicate that patient CASPR2 Abs enhance spontaneous network activity in both human and rodent neurons, providing a potential cellular basis for the network hyperexcitability associated with CASPR2-antibody encephalopathy and seizures.

**Fig. 1.**
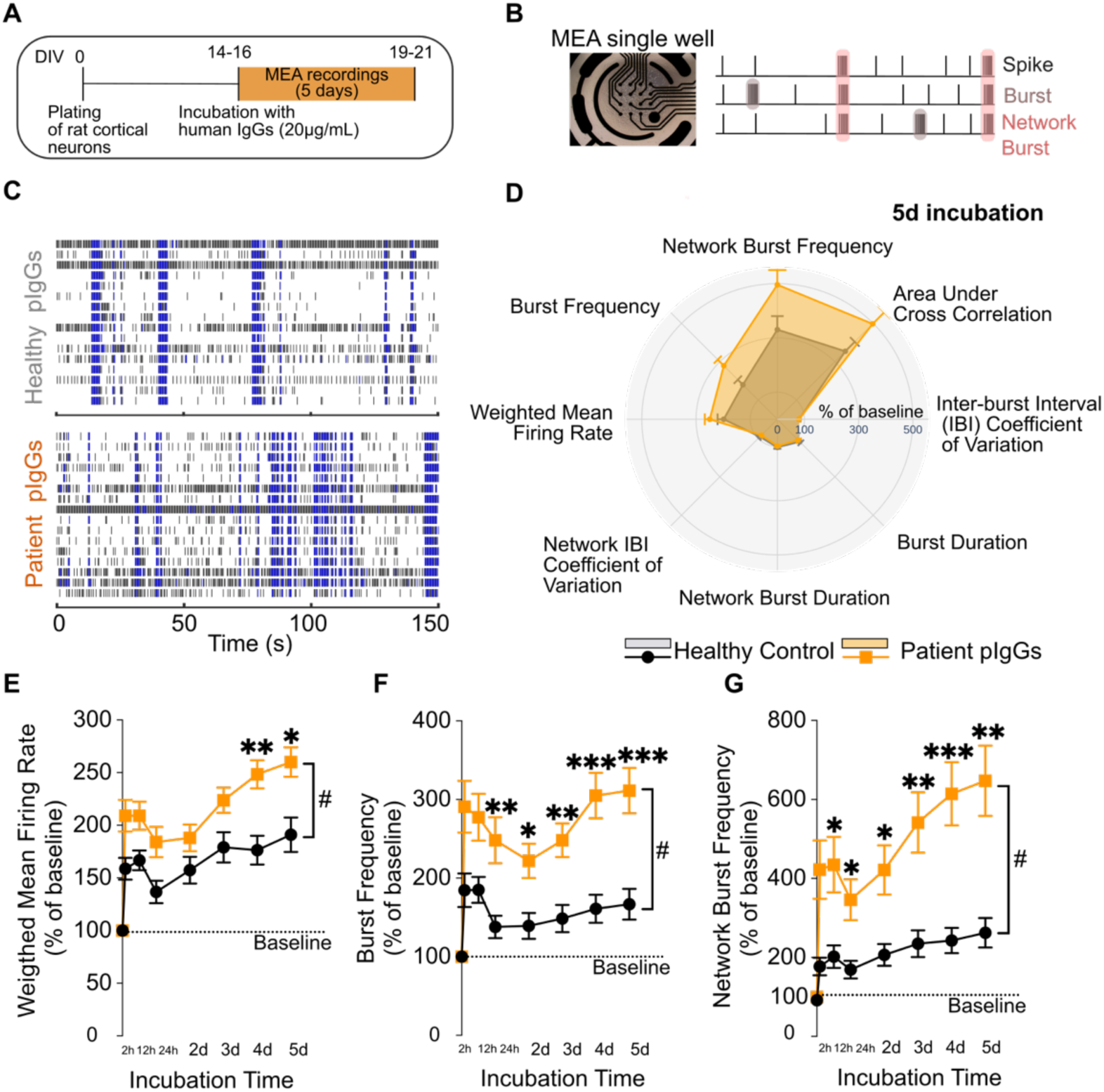
Patient-derived CASPR2 antibodies increase neuronal network activity in rat cortical neurons. (**A**) Schematic of the experimental design. (**B**) Schematic diagram of a MEA well and representative raster plot. (**C**) Representative 150-second raster plots of spontaneous activity of cultures at 5 days of incubation with the human pIgGs. Individual spikes are represented in black and spikes in bursts are in blue. (**D**) Quantification of the Weighted Mean Firing Rate, Burst Frequency and Duration, Network Burst Frequency and Duration, Inter-burst Interval (IBI) Coefficient of Variation, Network IBI Coefficient of Variation, and Area Under Cross-Correlation. The radar plot shows that Patient pIgGs increased network activity, reflected by increases in Weighted Mean Firing Rate, Burst Frequency, Network Burst Frequency, and Area Under Cross-Correlation at 5 days of incubation. Data are expressed as the percentage of baseline activity recorded immediately before the application of human pIgGs. (**E-G**) Patient pIgGs increased Weighted Mean Firing Rate (**E**), Burst Frequency (**F**), and Network Burst Frequency (**G**) compared to healthy control pIgGs over 5 days of incubation. Statistical analysis was performed on data shown in panels d–g (n = 37 wells from three healthy control pIgGs; n = 116 wells from Patient pIgGs from three individual patients, N=7 independent experiments), using a two-way Repeated Measures ANOVA test followed by Sidak’s multiple comparisons; *p<0.05, **p<0.01, ***p<0.001 indicate a significant *Time x pIgG treatment* difference compared to healthy control pIgGs; # p<0.05 indicates a significant main effect of pIgGs treatment. Results are presented as mean ± S.E.M. *Abbreviations:* pIgGs, purified Immunoglobulins G.

### CASPR2 antibodies enhance the synaptic excitatory drive onto neurons

To dissect the cellular mechanisms underlying the CASPR2-Ab-induced network hyperexcitability, including synaptic, intrinsic, and passive membrane properties of both excitatory and inhibitory neurons, we performed a comprehensive functional characterization of individual excitatory *versus* inhibitory neurons (identified by hDLX-driven mCherry expression) using whole-cell patch-clamp recordings (Fig. 2A).

**Fig. 2.**
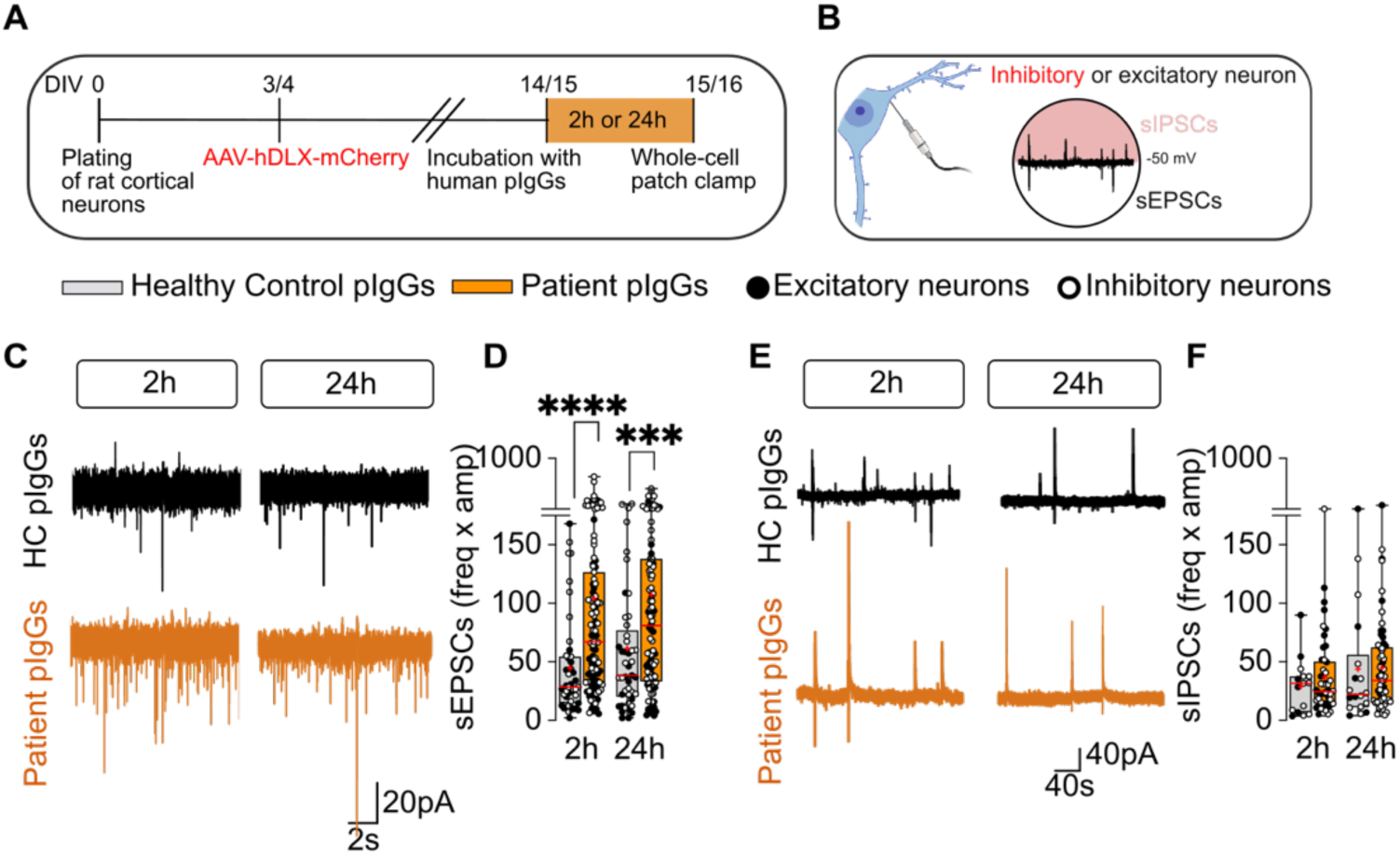
Patient-derived CASPR2 antibodies increase spontaneous excitatory transmission in rat cortical neurons. (**A,B**) Experimental design and patch-clamp protocols. (**C**) Representative traces showing the effects of human pIgGs on sEPSCs. (**D**) Quantification of sEPSCs, plotted as frequency (Freq) x amplitude (Amp) for each individual neuron. Patient pIgGs significantly increased excitatory drive (sEPSCs Freq x Amp) after both 2 h and 24 h of incubation. (**E**) Representative traces showing the effects of human pIgGs on sIPSCs. (**F**) Quantification of sIPSCs, plotted as Freq x Amp for individual neurons. Patient pIgGs did not alter inhibitory synaptic transmission at either 2 h or 24 h of incubation. For sEPSCs data at 2 h, N = 6 independent experiments [n(Healthy Control pIgGs) = 43 neurons, n(Patient pIgGs) = 111 neurons; 24 h, N = 5 independent experiments, n(Healthy Control pIgGs) = 37 neurons, n(Patient pIgGs) = 82 neurons]. For sIPSCs data at 2 h, N = 6 independent experiments [n(Healthy Control pIgGs) = 15 neurons, n(Patient pIgGs) = 46 neurons; 24 h, N = 5 independent experiments, n(Healthy Control pIgGs) = 17 neurons, n(Patient pIgGs) = 56 neurons]. Comparisons between conditions at 2 h or 24 h were performed using Mann–Whitney tests; ***p<0.001, ****p<0.0001 indicate a significant difference compared to healthy control <u>pIgGs</u>. Results in panels d and f are presented as boxes showing the 25^th^ and 75^th^ percentiles, with whiskers showing the range (minimum to maximum values corresponding to individual cells). The horizontal line within the boxes indicates the median. Crosses indicate the mean.

To assess spontaneous synaptic transmission, neurons were voltage-clamped at -50mV to record sEPSCs and sIPSCs simultaneously (Fig. 2B; see Methods, Electrophysiology section). Exposure to patient pIgGs for either 2h or 24h enhanced excitatory synaptic transmission (Fig. 2C,D), as indicated by the Frequency x Amplitude metric, calculated for each cell as the product of event frequency and mean amplitude of sEPSCs. At 2h, patient antibodies increased the Frequency x Amplitude of sEPSCs and amplitudes in both excitatory and inhibitory neurons, with increased sEPSC frequency in excitatory neurons (Fig. S4A-F). After 24h incubation, sEPSC frequency was only increased in inhibitory neurons (Fig. S4G-L). Notably, patient pIgGs did not alter inhibitory synaptic transmission (Fig. 2E,F, S4D-F, S4J-L), except for an increase in sIPSC amplitude in excitatory neurons following 24h incubation (Fig. S4J). Overall, these findings indicate that exposure to CASPR2 Abs predominantly enhances excitatory synaptic drive.

To determine whether patient-derived CASPR2 Abs alter neuronal membrane properties that influence intrinsic excitability, we measured membrane capacitance, input resistance, and resting membrane potential (RMP). Exposure to patient pIgGs did not significantly affect these properties in comparison to control pIgGs at either 2h or 24h of incubation (Fig. 3A-C). However, cell type-specific analyses showed that excitatory neurons exposed to patient pIgGs for 24h, presented a more depolarized RMP (Fig. S5) and reduced rheobase (Fig. 3D-F).

**Fig. 3.**
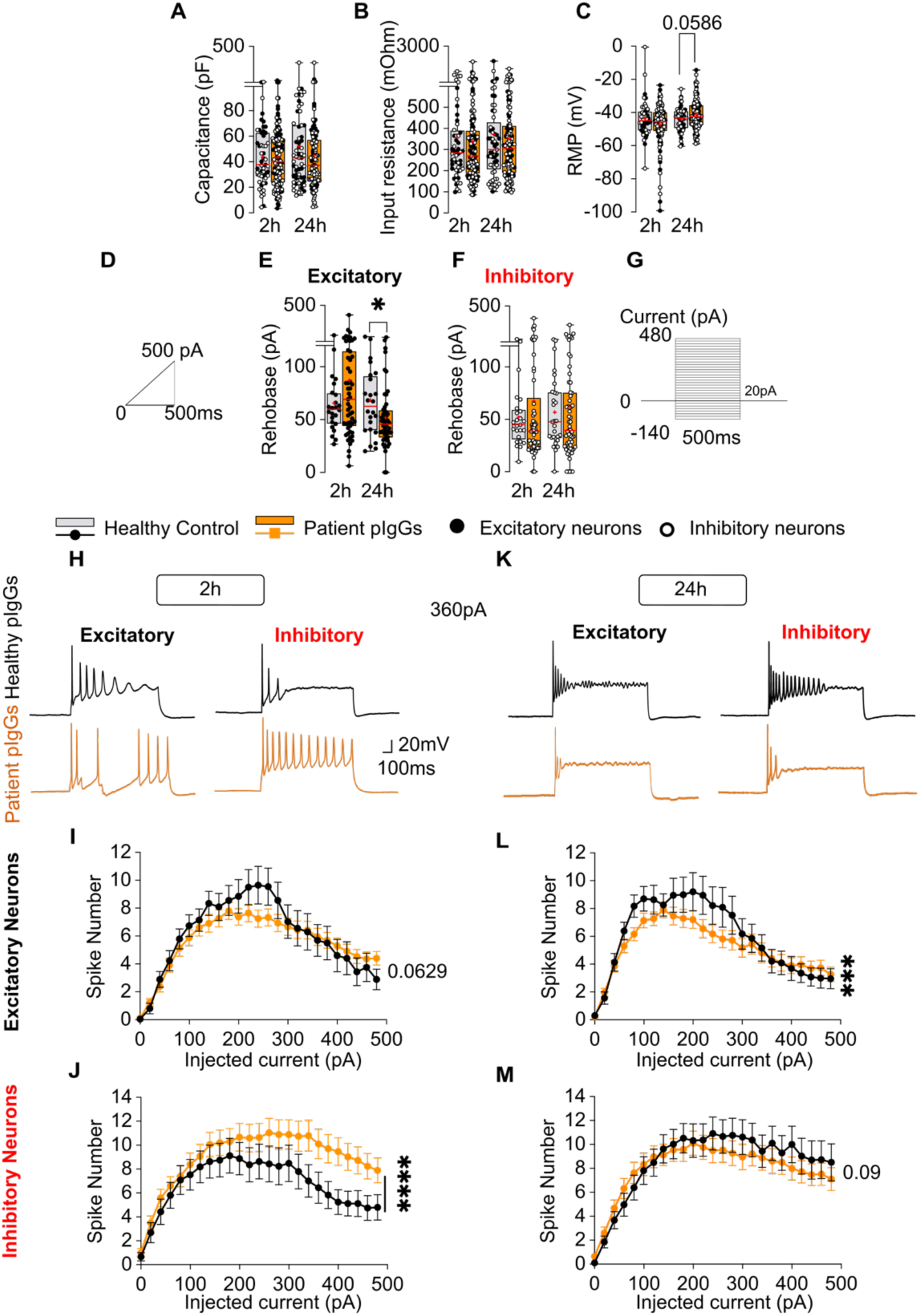
Patient-derived CASPR2 antibodies alter the intrinsic excitability of rat cortical neurons. Whole-cell patch-clamp recordings were performed following the experimental design in 2A. Purified IgGs from individual patients were tested separately and data were pooled as Patient pIgGs. (**A-C**) Passive membrane properties. (**A**) Membrane capacitance; (**B**) input resistance, determined from the steady-state current response to a 10 mV voltage step; and (**C**) resting membrane potential (RMP), measured at I = 0. Patient pIgGs did not significantly alter any of these parameters after either 2h or 24h. (**D**) Schematic of the current-ramp protocol used to determine rheobase (0–500 pA over 500 ms). (**E,F**) Rheobase of excitatory (**E**) and inhibitory (**F**) neurons. Patient pIgGs decreased rheobase in excitatory neurons after 24 h, but not after 2 h, with no significant effect in inhibitory neurons. (**G**) Schematic of the current-clamp protocol used to assess neuronal excitability. Neurons received 500-ms current injections in 20-pA increments from −140 to +480 pA. (**H,I**) Representative current-clamp traces after 2 h (**H**) and 24 h (**I**) of incubation with control or patient pIgGs, shown for excitatory (Left) and inhibitory (Right) neurons, and obtained from the +360 pA current step. J-M. Number of action potentials evoked at each current step. After 2 h, Patient pIgGs increased firing in inhibitory neurons but did not affect excitatory neurons (J,K). After 24 h, Patient pIgGs decreased firing in excitatory neurons but did not affect inhibitory neurons (L,M). For panels A-C, statistical comparisons were performed separately at each time point using the Mann–Whitney test, except for RMP, for which an unpaired t-test or Mann–Whitney test was used according to data distribution. For panels E,F, conditions were compared separately at each time point using the Mann–Whitney test. For panels J-L, data were analysed using two-way ANOVA; *p<0.05, ***p<0.001 and ****p<0.0001 indicate a significant main effect of pIgG treatment. Statistical analyses were based on six independent experiments per time point. The numbers of cells analysed were: a, 2 h: 51 control/130 patient; 24 h: 59/112; b, 2 h: 52/127; 24 h: 57/117; c, 2 h: 48/120; 24 h: 51/111; e, 2 h: 24/55; 24 h: 24/50; f, 2 h: 25/58; 24 h: 29/61; j, 2 h: 20/57; 24 h: 23/47; k, 2 h: 25/61; 24 h: 27/60 (Healthy Control pIgGs/Patient pIgGs, respectively). In panels A,B, boxes represent the 25^th^–75^th^ percentiles, whiskers the minimum–maximum range, the horizontal line the median, and crosses the mean. In panels J-M, results are represented as mean ± SEM. Unless otherwise indicated, differences were not statistically significant.

We next investigated the effects of CASPR2 Abs on neuronal firing by injecting square current steps of increasing amplitude (Fig. 3G). In response to suprathreshold current injection, inhibitory neurons showed an increased number of APs versus injected current after 2h of patient pIgG incubation (Fig. 3H-J), which was no longer observed after 24h (Fig. 3K-M). In contrast, excitatory neurons exhibited reduced firing following 24h of pIgG exposure (Fig. 3L). We found no major effects of patient pIgGs on AP kinetics (Fig. S6). Hence, CASPR2 Abs modulate neuronal firing in a cell type- and exposure time-dependent manner in response to the application of suprathreshold current injection. Yet, taken together, our results show CASPR2 Abs enhance excitatory synaptic drive, while selectively increasing the intrinsic excitability of excitatory neurons, as evidenced by changes in resting membrane potential and rheobase (which are measured under physiological current injections).

### The soluble CASPR2 ectodomain is an antigenic target of patient antibodies

We next investigated molecular mechanisms underlying these functional effects. CASPR2 is known to undergo activity-dependent ectodomain shedding mediated by matrix metalloproteinase 9 (MMP9), generating a soluble extracellular fragment detectable in human cerebrospinal fluid (CSF) and which plays a critical role in the homeostatic control of network neuronal activity (*27*). Therefore, we hypothesized that CASPR2 Abs could alter the levels and/or function of the soluble CASPR2 ectodomain and contribute to network hyperexcitability.

To test this hypothesis, we first assessed whether the shed CASPR2 ectodomain was present in the extracellular culture medium (ECM) of rat cortical neuronal cultures by immunoblotting after TCA precipitation (Fig. 4A) A distinct N-terminal CASPR2-immunoreactive band was detected in the ECM, migrating at a lower molecular weight than full-length CASPR2 present in whole-cell lysates (WCL; Fig. 4B,C). In contrast, the intracellular C-terminus of CASPR2 was detected exclusively in the WCL (Fig. 4C). The absence of actin in the ECM confirms that the media was not contaminated by cellular debris (Fig. 4C). These findings are consistent with the presence of the shed CASPR2 ectodomain in the ECM. To assess if patient CASPR2 antibodies could bind the soluble CASPR2 ectodomain, rat cortical neurons were incubated for 24h with patient- or healthy control-derived pIgGs (20 µg/mL; Fig. 4D), after which pIgGs were immunoprecipitated for analysis by immunoblotting. All three patient pIgGs, but none of the control pIgGs, immunoprecipitated the shed CASPR2 ectodomain, resulting in its depletion from the ECM (Fig. 4E-G).

**Fig. 4.**
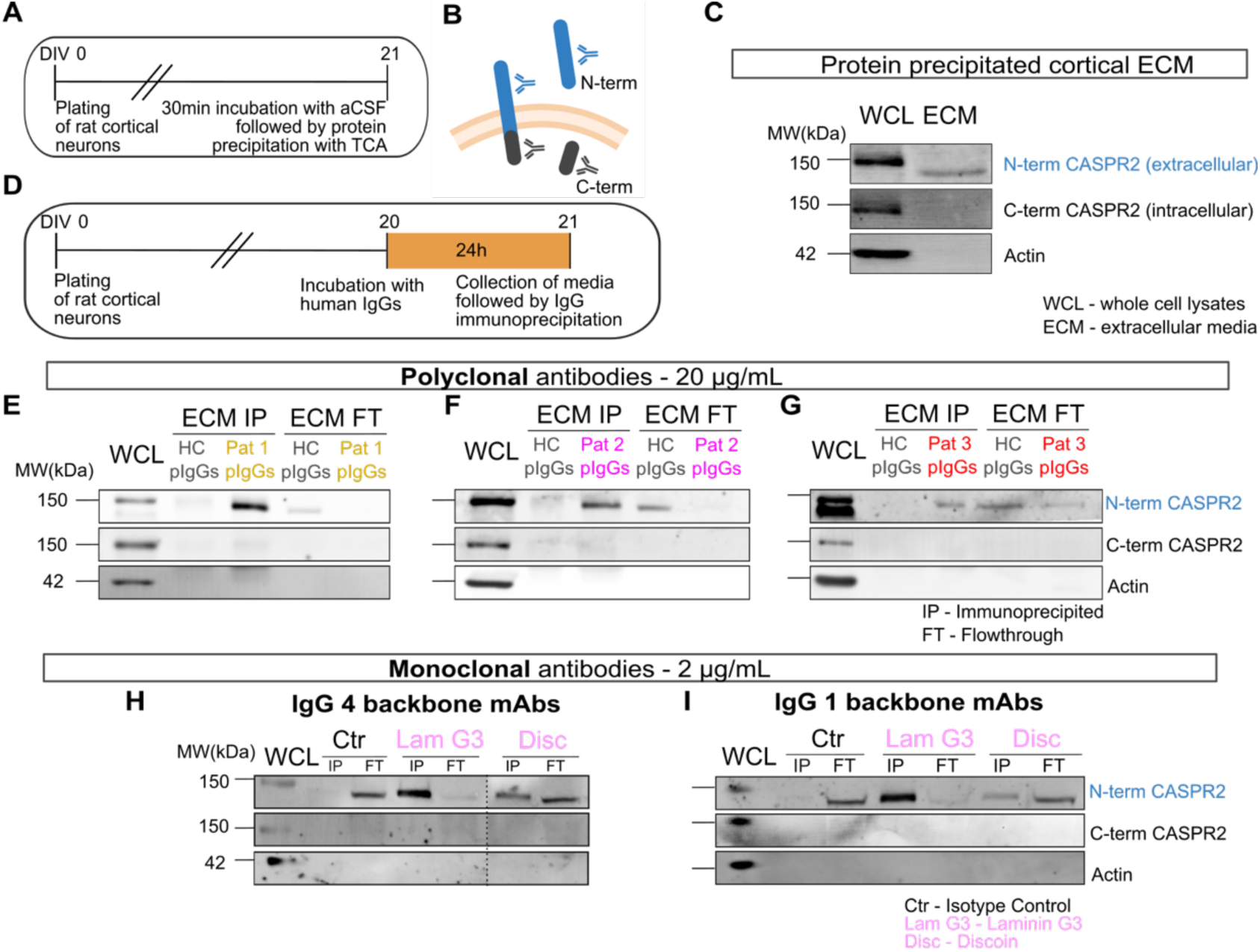
Patient-derived CASPR2 antibodies bind to the soluble CASPR2 ectodomain found in the extracellular medium of rat cultured neurons. **(A)** Experimental design for protein precipitation from the extracellular medium (ECM) of primary rat cortical cultures, using tricholoroacetic acid (TCA). **(B)** Illustration of commercial antibody epitopes on CASPR2, showing C-terminal (C-term, brown) and N-terminal (N-term, blue) antibody recognition sites. (**C**) Western blot (WB) analysis of the precipitated ECM proteins and whole-cell lysates (WCL) from the same culture, probed with commercial N-term and C-term CASPR2 antibodies. (**D**) Schematic of the experimental design for the immunoprecipitation of human polyclonal IgGs or recombinant monoclonal IgGs from the ECM. (**E-G**) Immunoblots showing that the CASPR2 pIgGs found in Patients 1 (**E**), 2 (**F**) and 3 (**G**) bound specifically to the soluble CASPR2 ectodomain (strong N-term band), depleting it from the culture medium (N-term band absent or strongly reduced in the flowthrough - FT). (**H,I**) Immunoblots showing that two monoclonal antibodies (mAb) targeting the laminin G3 and discoidin domains of CASPR2, produced recombinantly in (**H**) IgG4 and (**I**) IgG1 backbones, bound to shed CASPR2, whereas isotype controls did not.

To determine whether this ectodomain captured the majority of the extracellular domain of CASPR2, we next tested two CASPR2-anitbody encephalitis patient-derived monoclonal antibodies (mAbs) recognizing distinct extracellular domains of CASPR2 (*15*). Both mAbs, targeting the C-terminal laminin G3 domain and N-terminal discoidin domain, immunoprecipitated the CASPR2 ectodomain from the ECM (Fig. 4H,I). These findings indicate that epitopes found in both distal and proximal regions of the extracellular domain of CASPR2 are preserved in the soluble CASPR2 ectodomain and are available for patient antibody binding.

These results provide the first evidence that the shed CASPR2 ectodomain is a target of patient autoantibodies, and hence a previously unrecognized antigenic target in CASPR2-antibody diseases.

### CASPR2 antibodies impair calcium homeostasis in a PMCA2-dependent manner

The physiological effects of the soluble CASPR2 ectodomain on neuronal network activity have been reported to be mediated by the plasma membrane Ca^2+^-ATPase PMCA2, whereby the ectodomain constitutively binds PMCA2 and enhances its activity, promoting calcium extrusion to maintain neuronal calcium homeostasis (*27*). Hence, we concluded that physiological levels of the CASPR2 ectodomain were critical for neuronal homeostasis and that patient antibodies may alter neuronal excitability by disrupting intracellular calcium homeostasis through impaired CASPR2 ectodomain-mediated activation of PMCA2.

To test this hypothesis, we performed live-cell calcium imaging experiments in rat cortical neuron cultures using established protocols (*27, 33*). Neurons were co-transduced with AAVs expressing hDLX-mCherry, to identify inhibitory neurons, and synapsin-driven GCaMP6f, to monitor intracellular calcium dynamics (Fig. 5A). Following 24h incubation with individual control or patient pIgGs (20 µg/mL), neurons were depolarized with 30 mM KCl and changes in GCaMP6f fluorescence were recorded by live-imaging as a readout of intracellular calcium dynamics (Fig. 5B,C). In neurons incubated with patient pIgGs, depolarization elicited larger calcium responses, as measured by GCaMP6f fluorescence area under curve (AUC) (Fig. 5C-F, I, J), and altered calcium signal kinetics, reflected in an increased half-time of decay (t_1/2_off; Fig. 5G,H,K,L). These results confirmed that patient pIgGs impair calcium homeostasis, leading to slowed calcium clearance and prolonged calcium transients.

**Fig. 5.**
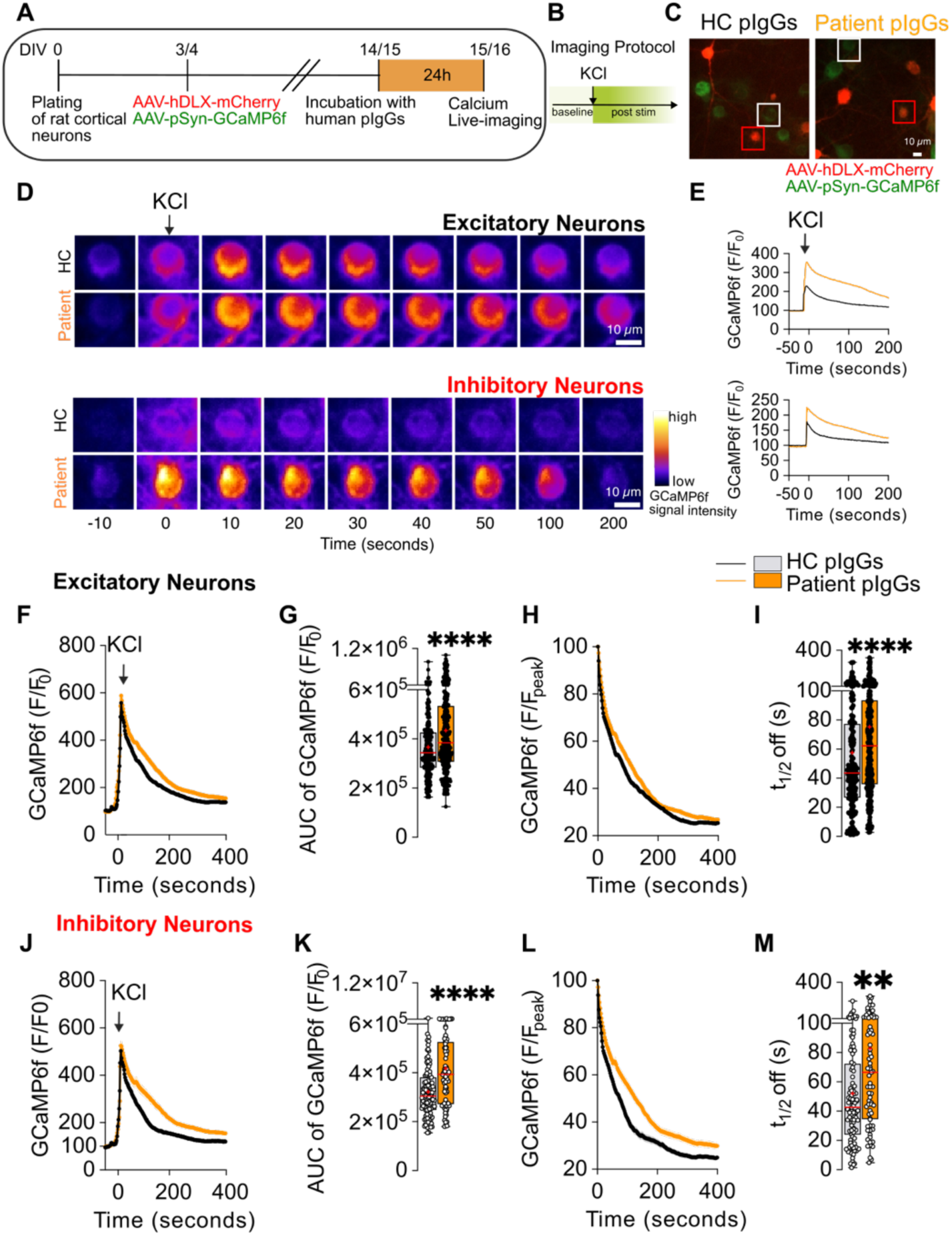
Patient-derived CASPR2 antibodies disrupt calcium homeostasis in rat cortical neurons. (**A**) Schematic of the experimental design. Neurons were incubated with pIgGs from each patient separately, but data from all patient samples were combined for analysis as Patient pIgGs. (**B**) Calcium dynamics in the somata of neurons in response to KCl-induced depolarization were assessed using live-cell imaging. (**C**) Epifluorescence images of transduced neurons preincubated with human pIgGs. Red insets indicate mCherry^+^ inhibitory neurons, and white insets indicate mCherry^-^excitatory neurons. (**D**) Magnification of the selected insets showing soma calcium signals in response to depolarization over time. (**E**) Representative traces corresponding to the average Ca^2+^ signals normalized to the baseline of the soma shown in D. (**F-I**) Calcium responses in excitatory neurons. (**F**) Average Ca^2+^ signals normalized to the baseline. (**G**) AUC of the individual responses. (**H**) Average Ca^2+^ signals normalized to the maximal amplitude. (**I**) t_1/2_ off values quantified from curves fitted to the decay phase of individual calcium signals. (**J-M**) Calcium responses in inhibitory neurons. (**J**) Average Ca^2+^ signals normalized to the baseline. (**K**) AUC. (**L**) Normalized decay. (**M**) t_1/2_ off values. Statistical analyses were performed on data from N = 6 independent experiments [For excitatory neurons, n(Healthy Control pIgGs) = 228 neurons, n(Patient pIgGs) = 299 neurons; For inhibitory neurons, n(Healthy Control pIgGs) = 103 neurons, n(Patient pIgGs) = 73 neurons]. AUC and t_1/2_ off values were analyzed using the Mann-Whitney test; **p < 0.01, ****p < 0.0001 indicates significance in comparison with Healthy Control pIgGs. Results in panels E,F,H,J and L are presented as mean ± S.E.M. Results in panels G,I,K and M are presented as boxes showing the 25^th^ and 75^th^ percentiles, with whiskers showing the range (minimum to maximum values corresponding to individual cells). The horizontal line within the boxes represents the median. Crosses indicate the mean. *Abbreviations:* pIgGs, purified Immunoglobulins G; AUC, Area Under Curve; HC, healthy control; F, Fluorescence; t_1/2_, decay half-time.

Next, we assessed whether these changes were PMCA2-dependent, by imaging calcium dynamics in neurons following PMCA2 knockdown (Fig. 6A). Neurons were co-transduced with AAV-GCaMP6f and lentiviruses expressing either PMCA2-targeting shRNA or a non-targeting scramble shRNA, both co-expressing mCherry (Fig. 6B; see Fig. S7 for validation of PMCA2 knockdown). Since excitatory and inhibitory neurons exhibited similar responses to KCl-induced depolarization and were similarly affected by patient IgGs (Fig. 5), we analyzed both populations together when assessing the effects of patient IgGs and PMCA2 depletion. As expected, in scramble-infected neurons, patient pIgGs increased the GCaMP6f signal AUC (Fig. 6E,F) and slowed calcium decay kinetics (Fig. 6G,H) compared to control pIgGs. However, these differences were lost in PMCA2 KD conditions (Fig. 6E,H), indicating that patient pIgGs impair calcium extrusion dynamics specifically through PMCA2. Overall, these findings demonstrate that CASPR2 antibodies disrupt calcium homeostasis in neurons through a mechanism dependent on PMCA2 activity.

**Fig. 6.**
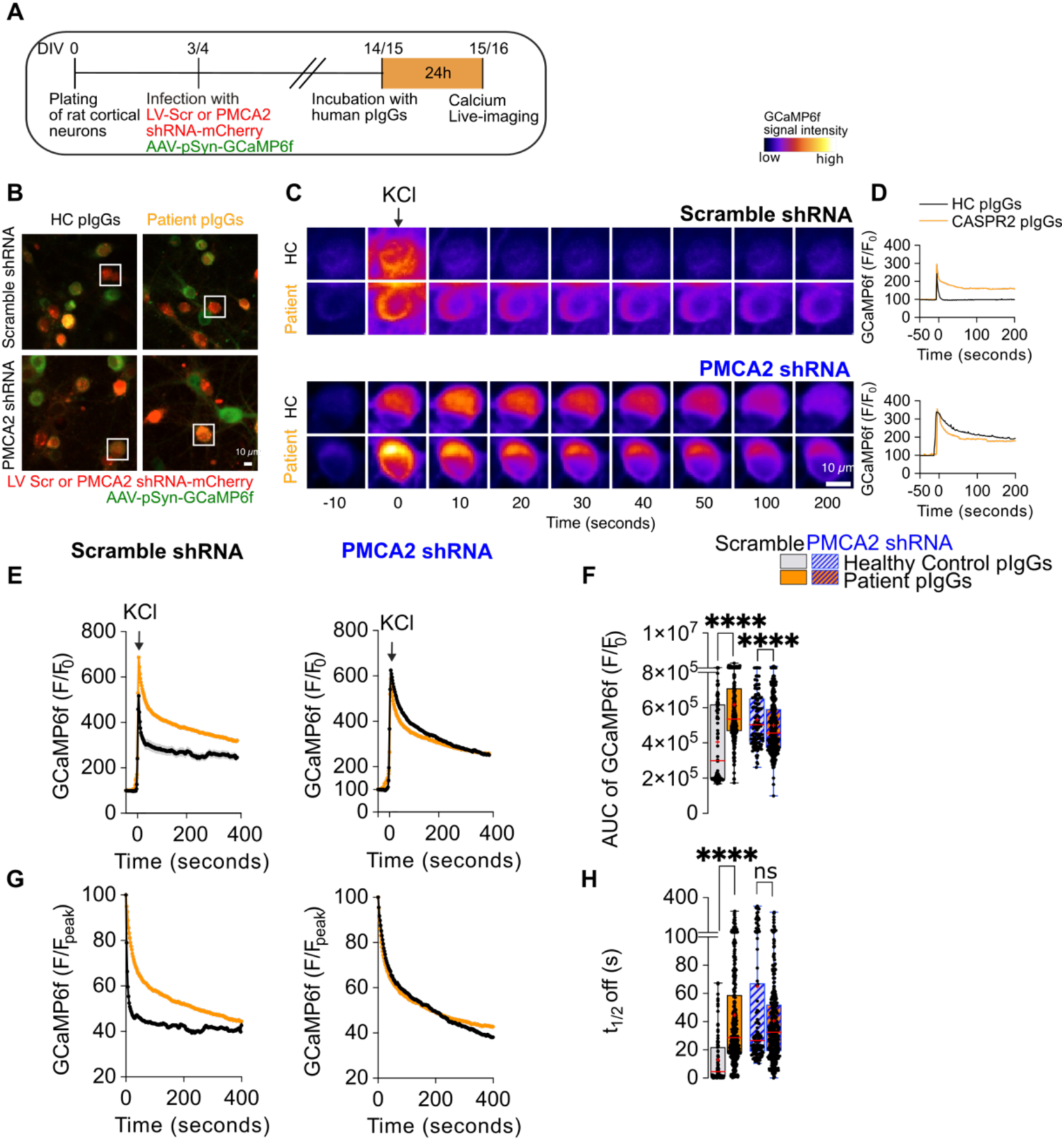
Patient-derived CASPR2 antibodies impair calcium clearance in a PMCA2-dependent manner. (**A**) Schematic of the experimental design. (**B**) Epifluorescence images of transduced neurons preincubated with human pIgGs, prior to KCl depolarization. Insets indicate mCherry^+^ neurons expressing shRNA constructs. (**C**) Time-course showing soma calcium signals in response to depolarization for the neuron in the insets in b. (**D**) Corresponding calcium response traces from somata of mCherry^+^ neurons under the four conditions. (**E-H**) Calcium responses in mCherry^+^ neurons. (**E**) Average traces of KCl-elicited Ca^2+^ signals in scramble shRNA (Left) or PMCA2 shRNA (Right) conditions. (**F**) AUC of the individual calcium responses. (**G**) Ca^2+^ decay phase normalized to maximal amplitude in scramble shRNA (Left) or PMCA2 shRNA (Right) conditions. (**H**) t1/2 off values of KCl-elicited Ca^2+^ peaks. Statistical analysis was performed on data from N = 3 independent experiments [n(Scramble shRNA, Healthy Control pIgGs) = 86 neurons, n(Scramble shRNA, Patient pIgGs) = 335 neurons, n(PMCA2 shRNA, Healthy Control pIgGs) = 118 neurons, n(PMCA2 shRNA, Patient pIgGs) = 339 neurons]. AUC and t_1/2_ off values were analyzed using a two-way ANOVA followed by Sidak’s multiple-comparisons test; ****p < 0.0001 indicates a significant difference compared to Healthy Control pIgGs. Results in panels e and g are presented as mean ± S.E.M. Results in panels f and h are presented as boxes showing the 25^th^ and 75^th^ percentiles, with whiskers showing the range (minimum to maximum values corresponding to individual cells). The horizontal line within the boxes represents the median. Crosses indicate the mean. Abbreviations: pIgGs, purified Immunoglobulins G; AUC, Area Under Curve; HC, healthy control; F, Fluorescence; t_1/2_, decay half-time.

### PMCA2 is required for CASPR2 autoantibody-induced synaptic and neuronal network hyperexcitability

To more directly test if CASPR2 autoantibody-induced synaptic and network dysfunction is mediated by PMCA2, we assessed spontaneous synaptic currents following PMCA2 knockdown. Neurons were co-transduced with lentiviruses expressing either non-targeting scramble shRNA or PMCA2-targeting shRNA and mCherry, and AAVs encoding mDLX-GFP to identify inhibitory (GFP+) and excitatory (GFP-) neurons. Following 24h exposure to control or patient pIgGs, whole-cell patch-clamp recordings were performed at a holding potential of –50 mV to record spontaneous post-synaptic currents (Fig. 7A). Consistent with our previous observations, in scramble control neurons, patient pIgGs increased sEPSCs compared to control pIgGs (Fig. 7B,C). This effect was not observed after PMCA2 knockdown (Fig. 7B,C), demonstrating that CASPR2 antibodies increase synaptic excitation in a PMCA2-dependent manner. Analysis of individual synaptic parameters showed that this effect was primarily driven by elevated sEPSC frequency, which was prevented by PMCA2 knockdown (Fig. S8). In contrast, patient pIgGs altered neuronal intrinsic properties and evoked firing rates regardless of PMCA2 expression levels (Fig. S9), indicating that antibody-mediated changes in excitability are independent of PMCA2.

**Fig. 7.**
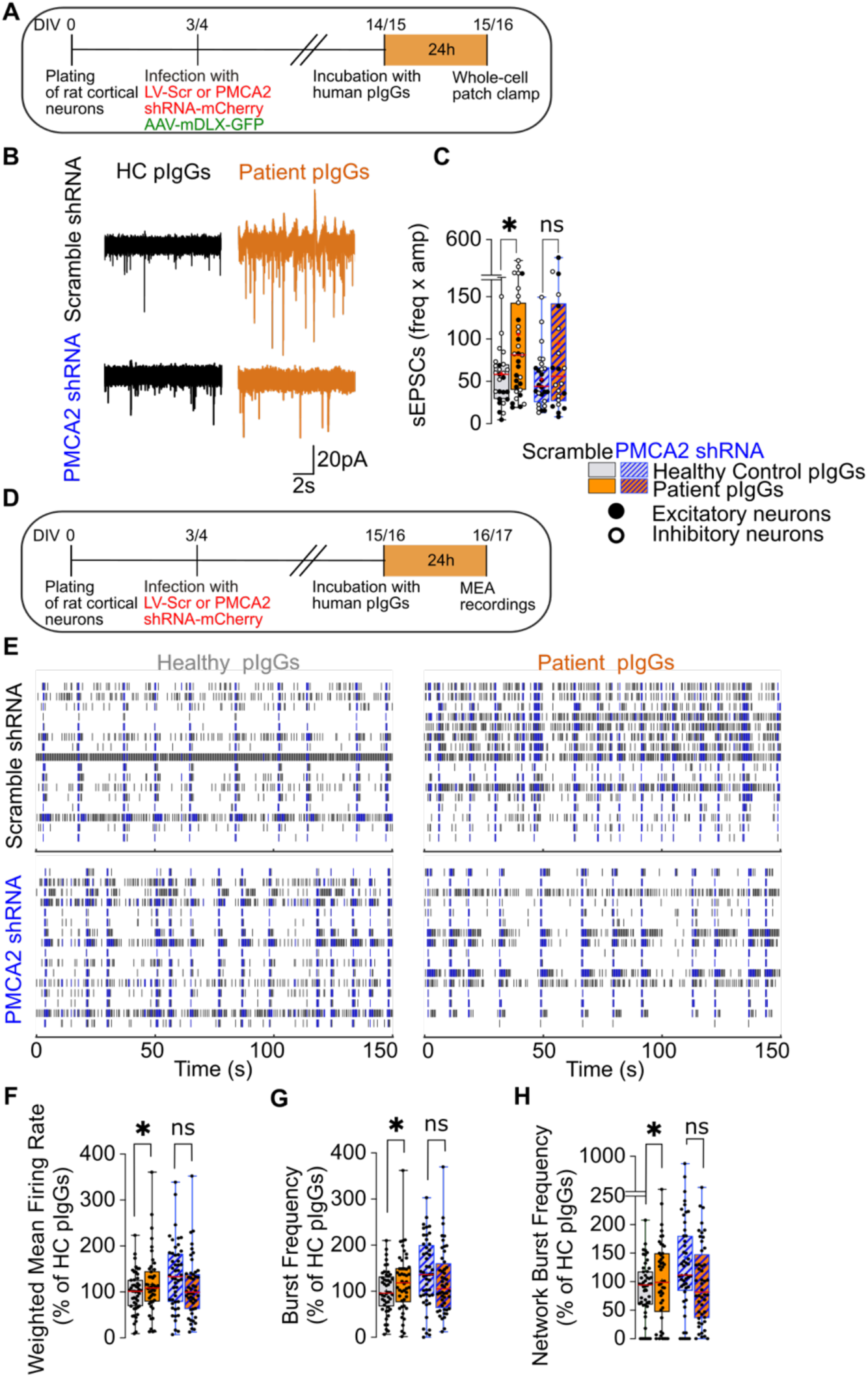
PMCA2 silencing prevents CASPR2 antibodies-induced increase in spontaneous excitatory synaptic transmission and network hyperexcitability in cortical neurons. **(A)** Schematic of the experimental design. **(B)** Representative traces showing the effects of human pIgGs on sEPSCs. **(C)** Quantification of the excitatory drive plotted as Freq x Amp for individual cells. Statistical analysis was performed on data from N = 5 independent experiments [n(Scramble HC pIgGs) = 29 neurons, n(Scramble Patient pIgGs) = 31 neurons, n(PMCA2 shRNA HC pIgGs) = 26 neurons, n(PMCA2 shRNA Patient pIgGs) = 22 neurons] using a two-way ANOVA followed by Sidak’s multiple-comparisons test. (**D**) Schematic of the experimental design to evaluate the role of PMCA2 in mediating the effects of CASPR2 autoantibodies on neuronal network function. In some experiments, pIgGs from individual patients were pooled prior to incubation. (**E**) Representative 150-second raster plots of spontaneous activity of cultures incubated with human pIgGs under the four experimental conditions. Individual spikes are represented in black, bursts are found in blue and network bursts are well-wide activity. (**F**) Weighted Mean Firing Rate, (**G**) Burst Frequency and (**H**) Network Burst Frequency. Data from individual wells were normalized to baseline and are shown as percentage of Healthy Control <u>pIgGs</u>. Statistical analyses were performed on data from N = 9 independent experiments [n(Scramble HC pIgGs) = 48 wells, n(Scramble Patient pIgGs) = 48 wells, n(PMCA2 shRNA HC pIgGs) = 46 wells, n(PMCA2 shRNA Patient pIgGs = 48 wells] using a two-way ANOVA followed by Sidák’s multiple comparisons post hoc test. Results in panel C,F,G and H are presented as boxes showing the 25^th^ and 75^th^ percentiles and whiskers showing the range (minimum to maximum values corresponding to individual wells). *p < 0.05 indicates a significant difference compared to Healthy Control pIgGs. The horizontal line within the boxes represents the median. Crosses indicate the mean. *Abbreviations:* HC, healthy control; pIgGs, purified Immunoglobulins G; LV, Lentiviruses; Freq, Frequency; Amp, Amplitude.

To test whether single-cell changes translate to network level dysfunction, we performed microelectrode array recordings in neurons transduced with lentiviruses expressing shRNA constructs following 24h incubation with human pIgGs (Fig. 7D). In Scramble shRNA conditions, patient pIgGs significantly increased spontaneous network activity, manifested as elevated Mean Firing Rate, Burst Frequency, and Network Burst Frequency, compared to healthy control pIgGs (Fig. 7E-H). These effects were abolished when PMCA2 expression was knocked down. Hence, the enhancement of excitatory synaptic transmission (Fig. 7A-C) and network activity (Fig. 7D-H) both require intact PMCA2 expression, whereas changes in intrinsic neuronal excitability persist independently of PMCA2 (Fig. S9).

This dissociation suggests that CASPR2 antibodies primarily disrupt network function through impaired calcium extrusion at the synaptic level, with secondary effects on intrinsic neuronal properties. Combined with the PMCA2-dependent disruption of intracellular calcium dynamics, these results support a model in which CASPR2 autoantibodies drive network hyperexcitability by impairing PMCA2-mediated calcium extrusion. In contrast, parallel PMCA2-independent mechanisms contribute to alterations in intrinsic excitability, likely involving previously described autoantibody targeting of full-length membrane CASPR2 and its interaction with ion channel complexes.

### CASPR2 antibodies in patient CSF bind CASPR2 ectodomain and reduce its free levels

To determine whether the pathogenic mechanism identified in neuronal cultures operates in patients, we investigated whether CASPR2 autoantibodies engage shed CASPR2 in the CSF. Samples from patients with high-titre CASPR2 antibodies and healthy controls were analyzed. Immunoprecipitation of CSF IgGs followed by Western blot revealed that patient autoantibodies were bound to the shed CASPR2 ectodomain in the CSF, while unbound levels of shed CASPR2 were markedly reduced in patient samples compared with controls (Fig. 8A-C), consistent with *in vivo* targeting of soluble CASPR2. To quantify this effect, shed CASPR2 levels were measured by ELISA before and after IgG immunoprecipitation (Fig. 8D,E). CASPR2 antibody-positive CSF contained substantially lower levels of shed CASPR2 both before and after immunoprecipitation, indicating depletion of the soluble ectodomain in patients. Together, these findings demonstrate that patient autoantibodies target and deplete physiologically shed CASPR2 in the CSF, providing clinical evidence that antibody-mediated *in vivo* engagement of a biologically active soluble ectodomain contributes to the pathogenesis of CASPR2 autoimmunity (See our current working model on Fig. S10).

**Fig. 8.**
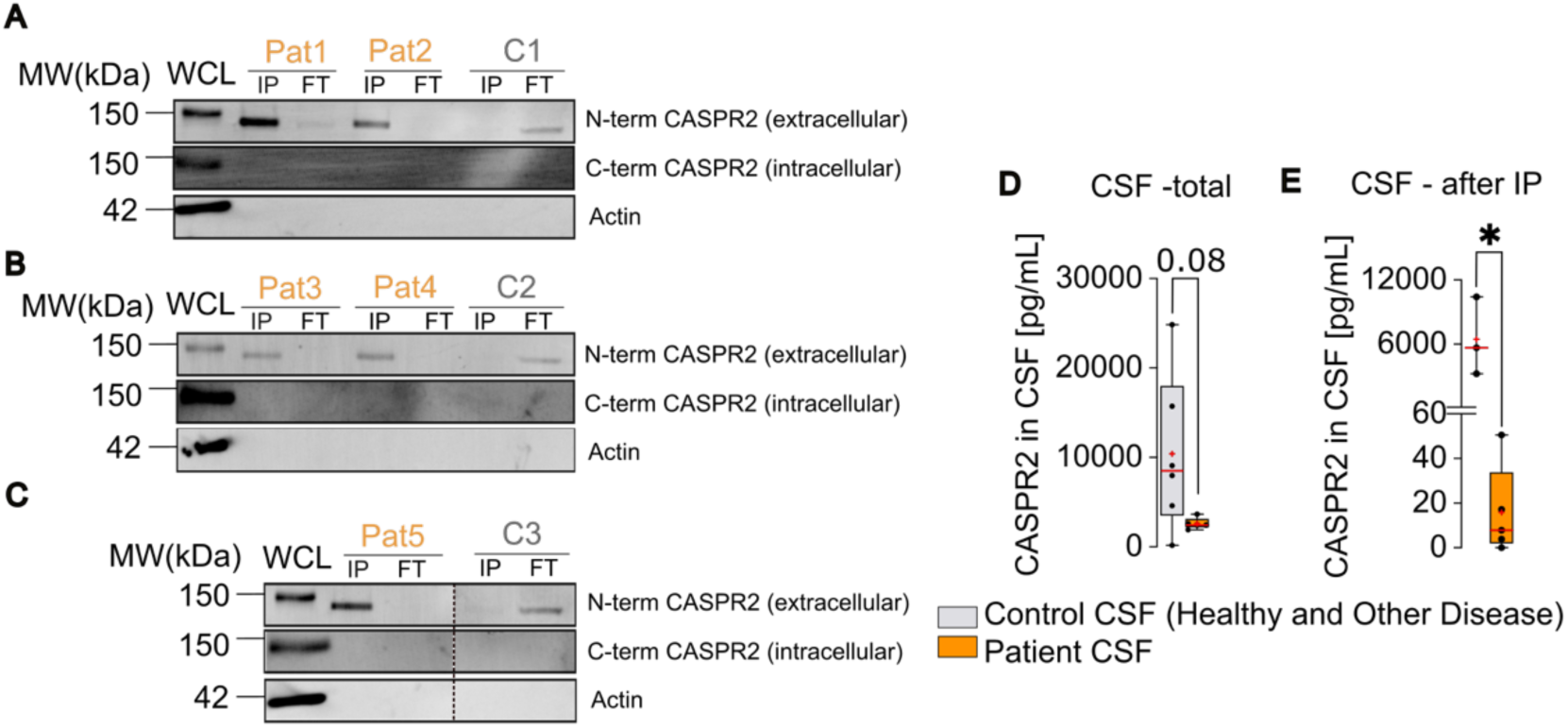
CASPR2 antibodies in the CSF of patients bind to shed CASPR2 and reduce its free levels in the CSF of limbic encephalitis patients. (A-C) Patient antibodies were immunoprecipitated from the CSF of 6 individual patients and 3 control individuals. The whole cell lysate (WCL) from rat cortical neurons was run as a positive control for the full-length CASPR2. Immunoprecipitation (IP) and flow-through (FT) samples were probed with commercial antibodies against the N-terminus and C-terminus of CASPR2 and with an antibody against β-actin. **(D,E)** Concentration of shed CASPR2 in human CSF from control and patient samples, before **(D)** and after **(E)** immunoprecipitation to remove antibody-bound CASPR2 from the CSF. Statistical analysis was performed in the data from 3-6 Control CSF samples and 5 Patient CSF samples, using the Mann-Whitney test. *p<0.05 compared to Control. Results in panels d and e are presented as boxes showing the 25^th^ and 75^th^ percentiles, whiskers show the range (minimum to maximum values corresponding to individual samples). The horizontal line within the boxes represents the median. Crosses indicate the mean. *Abbreviations:* C, Control; CSF, Cerebrospinal fluid.

## DISCUSSION

Here, we identify the physiologically active shed CASPR2 ectodomain as a previously unrecognized pathogenic target in CASPR2 autoantibody-mediated neurological diseases. Patient-derived CASPR2 Abs bind shed CASPR2 and reduce its availability, thereby impairing its function in regulating neuronal calcium homeostasis (Fig. S10). This leads to aberrant PMCA2-dependent calcium accumulation, enhanced excitatory synaptic transmission and network hyperexcitability. Collectively, these findings reveal a previously unrecognized disease mechanism for CASPR2 autoantibody-associated diseases, particularly in patients presenting with neuronal hyperexcitability, and establish a paradigm by which pathogenic antibodies can interfere with the function of soluble proteins.

Our data show that patient-derived CASPR2 Abs disrupt calcium homeostasis and selectively enhance excitatory synaptic transmission, while leaving inhibitory synaptic transmission unaffected, in both excitatory and inhibitory neurons. Additionally, patient-derived CASPR2 Abs drive cultured rat neuronal networks into a hyperconnected state. These *in vitro* findings provide a mechanistic link to the clinical phenotype observed in patients with CASPR2 autoimmune encephalitis (*1, 3, 4, 8, 9*), in which seizures are prominent and hyperkinetic manifestations, including myoclonus and tremor, are also observed (*4, 8, 9, 18, 34*). Consistent with our findings, EEG abnormalities and enhanced functional connectivity have been reported in human CASPR2-antibody encephalitis patients and in animals administered CASPR2 mAbs (*12, 35*). Similarly, prolonged exposure of unguided human neural organoids to CASPR2 antibodies induces a hyperexcitability phenotype (*36*). Further support comes from seizure-like phenotypes observed in *Cntnap2* mutant animals (*37*) and hyperexcitability found in cortical excitatory networks derived from iPSCs carrying *CNTNAP2* null mutations (*38*). Collectively, these findings indicate that that disruption of CASPR2 function, including signaling mediated by its shed ectodomain, is sufficient to induce network hyperexcitability. The network phenotype we observed resembles that induced by epileptogenic drugs such as pilocarpine (*39, 40*) and by patient-derived NMDAR antibodies (*41*), suggesting that distinct molecular mechanisms underlying seizure may converge on common phenotypes at the network level.

Our results further reveal that patient-derived CASPR2 Abs bind and sequester the shed CASPR2 ectodomain in patient CSF which identifies the shed CASPR2 ectodomain as a pathogenic target in CASPR2 autoimmunity. Although autoantibody targeting of shed proteins has been described before for collagen XVII/BP180 in blistering skin diseases (*42*), this mechanism has not previously been reported in neurological disorders. Our findings extend the pathogenic mechanisms of neuronal autoantibodies beyond the modulation of cell-surface proteins and suggest that antibody-mediated sequestration of soluble protein fragments may represent an important mechanism in autoimmune diseases.

Our study establishes the CSF sheddome as a previously underappreciated source of potential autoantigen targets in neurological disease. Proteolytic cleavage of neuronal surface proteins can generate soluble ectodomains with biological activities distinct from those of their membrane-bound precursors. Because they freely diffuse through the extracellular space, these fragments may be particularly susceptible to targeting by pathogenic autoantibodies that disrupt their signaling pathways. Profiling of the CSF sheddome combined with screening for patient antibody reactivity can reveal additional soluble antigens relevant in neurological autoimmunity.

The shed CASPR2 ectodomain was recently identified as an endogenous ligand for PMCA2 that regulates intracellular calcium homeostasis to maintain neuronal network activity within the physiological range (*27*). This prompted us to investigate whether the functional effects of CASPR2 Abs are mediated at least partially through disruption of shed CASPR2 signaling. Indeed, we found that patient-derived CASPR2 Abs bind to and reduce the levels of free shed CASPR2, an effect associated with impaired PMCA2-dependent intracellular calcium homeostasis. Consistent with this mechanism, the effects of CASPR2 Abs on calcium dynamics in response to a depolarizing stimulus, spontaneous excitatory neurotransmission and neuronal network activity were abolished by PMCA2 knockdown, demonstrating that these effects require intact PMCA2-mediated regulation of calcium homeostasis. In support of our findings, PMCA2 has already been implicated in seizure pathology, as it is downregulated following 24h of kainic acid–induced seizures (*43*) and its pharmacological inhibition selectively enhances excitatory, but not inhibitory, synaptic transmission (*44*).

Notably, our findings highlight the mechanistic heterogeneity of CASPR2 antibody-mediated effects. While effects on calcium dynamics, excitatory synaptic transmission and neuronal network activity require PMCA2, changes in intrinsic neuronal excitability persisted after PMCA2 knockdown. This suggests that CASPR2 Abs trigger at least two pathogenic pathways, which are not mutually exclusive. The first one is dependent on the targeting the shed CASPR2 ectodomain and its modulation of PMCA2 activity that affects calcium homeostasis and downstream pathways. The second one is independent of PMCA2 and impacts neuronal intrinsic properties and excitability, likely implicating the direct targeting of membrane CASPR2 and disrupting its regulation of voltage-gated Kv-potassium channels, as previously reported (*13, 14, 45*).

Altered calcium signaling has been reported in several models of disease associated with CASPR2 dysfunction, including those using CASPR2 Abs and *CNTNAP2* knockout models (*14, 30, 46, 47*). Here, we show that patient CASPR2 Abs primarily disrupt neuronal calcium homeostasis by targeting shed CASPR2, and that their effect is mediated by PMCA2.

Remarkably, shed CASPR2 levels were drastically reduced in patient CSF. Given that shed CASPR2 is required for proper neuronal function, this marked depletion suggests that its physiological roles are compromised in patients. Together with our *in vitro* findings showing that CASPR2 Abs disrupt the shed CASPR2–PMCA2 pathway and impair calcium homeostasis, these data support the possibility that such mechanism contributes to disease pathogenesis in CASPR2 autoimmune encephalitis.

Despite these findings, several limitations should be acknowledged. First, the functional effects of CASPR2 autoantibodies were primarily investigated *in vitro* using rodent and human-derived iPSCs neuronal cultures, which may not fully recapitulate the complexity of the human disease. Further studies using *in vivo* systems will be important to establish the relevance of the identified mechanism. Second, although our findings demonstrate that PMCA2 is required for several antibody-mediated functional effects, the contribution of other molecular interactors of the shed CASPR2 ectodomain remains unexplored. Third, the reduced levels of soluble CASPR2 detected in patient CSF warrant further investigation in larger patient cohorts to establish their clinical relevance and determine their relationship with disease severity, activity and outcome. Finally, the mechanisms underlying the reduction in CSF soluble CASPR2 remain to be elucidated, as antibody-mediated sequestration and altered ectodomain shedding may both contribute to its depletion. Future studies measuring CASPR2 shedding in the presence of CASPR2 Abs will be important to evaluate these possibilities. Irrespective of its origin, the bioavailable pool of free shed CASPR2 is depleted in disease, compromising the signaling pathways it mediates.

In conclusion, our findings provide the playground for several important clinical implications for diagnosis, disease stratification and treatment of diseases mediated by CASPR2 autoantibodies. Diagnostic protocols could be improved by assessing antibody binding to the soluble CASPR2 ectodomain, as a complement to currently available antibody screening approaches relying on LCBAs that detect antibodies against full-length, membrane-bound CASPR2. Moreover, because patient-derived CASPR2 Abs reduce the levels of the free shed CASPR2 ectodomain, quantifying soluble CASPR2 in patient plasma and CSF could provide a potential biomarker reflecting the disruption of this physiological signaling pathway.

Our data also identify potential therapeutic opportunities beyond immunotherapy. If depletion of shed CASPR2 significantly contributes to disease pathogenesis, strategies aimed at normalizing soluble CASPR2 in the CSF or restoring PMCA2 activity may successfully rescue neuronal dysfunction in CASPR2 autoimmunity and open new avenues for mechanism-based therapeutic interventions.

## MATERIALS AND METHODS

### Study design

This study aimed to investigate the effects of human CASPR2 autoantibodies on neuronal function and determine whether they target and interfere with the physiological functions of the shed CASPR2 ectodomain. We first characterized the functional effects of patient-derived antibodies *in vitro* using electrophysiology and calcium imaging assays. We then performed biochemical assays to establish whether shed CASPR2 is an antigenic target of these antibodies and whether antibody binding alters its extracellular availability. Given the role of shed CASPR2 in regulating neuronal activity through PMCA2, we next investigated whether the effects of shed CASPR2 depletion on neuronal calcium homeostasis, intrinsic excitability, synaptic transmission, and network activity were dependent on PMCA2 by knocking down its expression in neurons. Finally, we measured soluble CASPR2 levels in the CSF of patients to explore its potential as a disease biomarker and therapeutic target.

### Patient polyclonal antibody samples, recombinant monoclonal antibodies and commercial antibodies

All the samples were collected with informed consent from CASPR2 patients, healthy individuals, or patients with unrelated diseases (Mayo Clinic Jacksonville: IRB 23-010102 and University of Oxford: REC16/YH/0013). Immunoglobulin G was purified from the plasma and their features are listed in Table S1. Sera samples information are listed in Table S2. Information on CSF samples is detailed in Table S3. Human IgG was purified from plasma samples using Protein G Sepharose 4 Fast Flow resin (Cytiva, USA: 17061802). CASPR2-specific monoclonal antibodies (mAbs) were recombinantly generated from patient-derived B cells, expressed in IgG1 or IgG4 backbones and recognised distinct epitopes on CASPR2 (*15*). A non-binding control mAb (G2) purified from patient-derived B cells, and a recombinant mAb engineered to bind the gastrointestinal glycoprotein A33 (non-neuronally expressed) were used as negative controls (*15*). The commercial antibodies used in this study are listed on Table S4 on the supplementary materials.

### Rat primary neuronal cultures and neuronal transduction

Neuronal primary cultures were obtained from the hippocampi or cortices isolated from E17-18 Wistar rat embryos, Animal procedures were reviewed and approved by the responsible authorities, Direção Geral de Veterinária (DGAV), Portugal (licences 2019/024628 and 2025/47110/25-S). Neuronal cultures were prepared as previously described (*17, 48*) and cells were plated at different densities depending on the specific experiment to be performed (see supplementary material for details). Neurons were transduced at DIV3-4 using lentiviruses to KD PMCA2 expression at a MOI of 5. For the neuronal transduction using GCaMP6f and hDLX AAVs, infections were performed at DIV3 or 4 using dilutions adjusted for each viral preparation, typically between 1:500 and 1:750 in culture medium. Infection was confirmed visually by the expression of GFP or mCherry.

### Live calcium imaging with GCaMP6f

GCaMP6f-expressing neurons were incubated with human CASPR2 IgGs (20 µg/mL) at DIV14–15 for 24 h. Neurons were then imaged in aCSF using a Zeiss Cell Observer Spinning Disk microscope at 3 frames/s. Following a 60-s baseline recording, 30 mM KCl was applied to induce neuronal depolarization and cytosolic calcium responses (*27*). Fluorescence signals were analysed in FIJI by measuring somatic ROIs, with peak amplitudes and calcium decay kinetics quantified using GraphPad Prism.

### Precipitation of extracellular proteins

To detect endogenous soluble CASPR2 ectodomain, conditioned artificial cerebrospinal fluid (aCSF) from DIV21 cortical neuronal cultures was collected following a 30 min incubation. Extracellular proteins were precipitated using trichloroacetic acid (TCA) as previously described (*27*). See supplementary material for protocol details.

### Immunoprecipitation of soluble CASPR2 ectodomain from culture media or from CSF

To assess the interaction between patient IgGs and soluble CASPR2 ectodomain, human IgGs from patients or healthy controls (20 µg/mL) were incubated with DIV20 cortical neuronal cultures for 24 h. Conditioned media were subsequently collected and incubated overnight with Protein A Sepharose beads to immunoprecipitate human IgGs and associated proteins. Immunoprecipitatin was also used to investigate the presence of soluble CASPR2 ectodomain in human CSF samples and its association with patient IgGs. Following washing, immunoprecipitated proteins were eluted in Laemmli sample buffer. See supplementary material for protocol details.

### Enzyme-Linked Immunosorbent Assay (ELISA)

The concentration of CASPR2 ectodomain in cerebrospinal spinal (CSF) samples (diluted 4-fold) was determined using a commercial ELISA Kit (RayBiotech: #ELH-CASPR2-1) following the manufacturer’s instructions.

### Electrophysiology

Whole-cell patch-clamp recordings were performed in DIV15–16 hDLX-mCherry+ (inhibitory) and hDLX-mCherry− (excitatory) neurons following 2 or 24h incubation with human IgGs. Intrinsic excitability was assessed in current-clamp mode by measuring AP firing, rheobase and membrane properties. Spontaneous excitatory and inhibitory postsynaptic currents (sEPSCs and sIPSCs) were recorded in voltage-clamp mode at −50 mV. Signals were acquired using a HEKA EPC10 amplifier and analysed with Igor Pro and Clampfit software. See supplementary material for details.

### Microelectrode array (MEA) assays

For experiments with rat neurons, the assays were conducted according to previously published works (*50*). Spontaneous activity was recorded using the Maestro Edge system (Axion Biosystems) following incubation with patient or healthy control IgGs. Baseline activity was recorded for 10 min, followed by recordings at multiple timepoints after IgG exposure. Spike and burst activity were analysed using Axis Navigator and NeuralMetrics software. Data were normalized to baseline activity and, for PMCA2 knockdown experiments, additionally to healthy control IgG conditions. See supplementary material for details. For experiments with iNeurons, human iPSC-derived neurons were differentiated and plated onto the MEA plates. Spontaneous activity was recorded weekly from two weeks after differentiation using a MED64 Presto system (Alpha Med Scientific). At seven weeks, neurons were incubated with healthy control or CASPR2-positive patient sera (1:100 dilution), and activity was recorded for 10 min every hour over 6h. Spike and burst activity were analysed using MEA Symphony and Burst Analysis Table software. Wells with fewer than eight active electrodes were excluded. See supplementary material for details.

### Statistical analyses

For each experiment, the normality of the data was assessed by comparison with a theoretical normal distribution using the Shapiro-Wilk normality test. According to this, the statistical significance between groups was assessed either by parametric or non-parametric tests, according to their data distribution. Analyses were performed using GraphPad Prism 9.0 (GraphPad software, USA). For all tests, p<0.05 was considered statistically significant.

## Acknowledgments

We would like to thank Jeannette Schmidt for help in viral production and Luisa Cortes and the MICC team for assistance with microscopy imaging. Schematic images were created using Biorender.com. We thank all Synapse Biology laboratory members for technical assistance and for the indispensable discussion of the work.

## Funding

This work was supported by:

Fundacio la Marato de TV3 (to ALC), under the project 202202-30-31;

La Caixa Foundation/FCT (i4b) initiative (to PP) under the project LCF/PR/HP21/52310010;

FCT (to ALC) under project POCI-01-0145-FEDER-029452.

Institutional funding to CIBB-Center for Innovative Biomedicine and Biotechnology was provided by the European Regional Development Fund (ERDF) through the COMPETE 2020 — Operational Programme for Competitiveness and Internationalisation and by the EU Recovery and Resilience Facility and Portuguese national funds via FCT under the projects LA/P/0058/2020 [DOI:10.54499/LA/P/0058/2020], UID/PRR/4539/2025 [DOI: 10.54499/UID/PRR/04539/2025] and UID/04539/2025.

DS was supported by SFRH/BD/132008/2017 and the COVID-19 COVID/BD/151955/2021 fellowships, funded via FCT.

SRI is supported by:

the Wellcome Trust (104079/Z/14/Z);

the Medical Research Council (MRC) (MR/V007173/1);

the National Institute for Health Research (NIHR) Oxford Biomedical Research Centre (BRC);

the Mayo Clinic Robert and Arlene Kogod Center on Aging [Aging Nervous System, 2024].

UCB funding supported part of this study.

## Author contributions

Conceptualization: ALC, DS

Methodology: ALC, DS, EC, PP, LR. DF

Investigation: DS, EC, NV, VB, QTN, TK, EW, RW, SRI

Visualization: DS, NV

Funding acquisition: ALC, SRI

Project administration: ALC

Supervision: ALC

Writing – original draft: ALC, DS

Writing – review & editing: ALC, DS, EC, PP, LR, DF, NV, VB, QTN, TK, EW, RW, SRI

## Data and materials availability

All data present in the main text or the supplementary materials upon request.

