## Supplementary Materials for "Pathogenic human autoantibodies target the soluble CASPR2 ectodomain to induce network hyperexcitability"

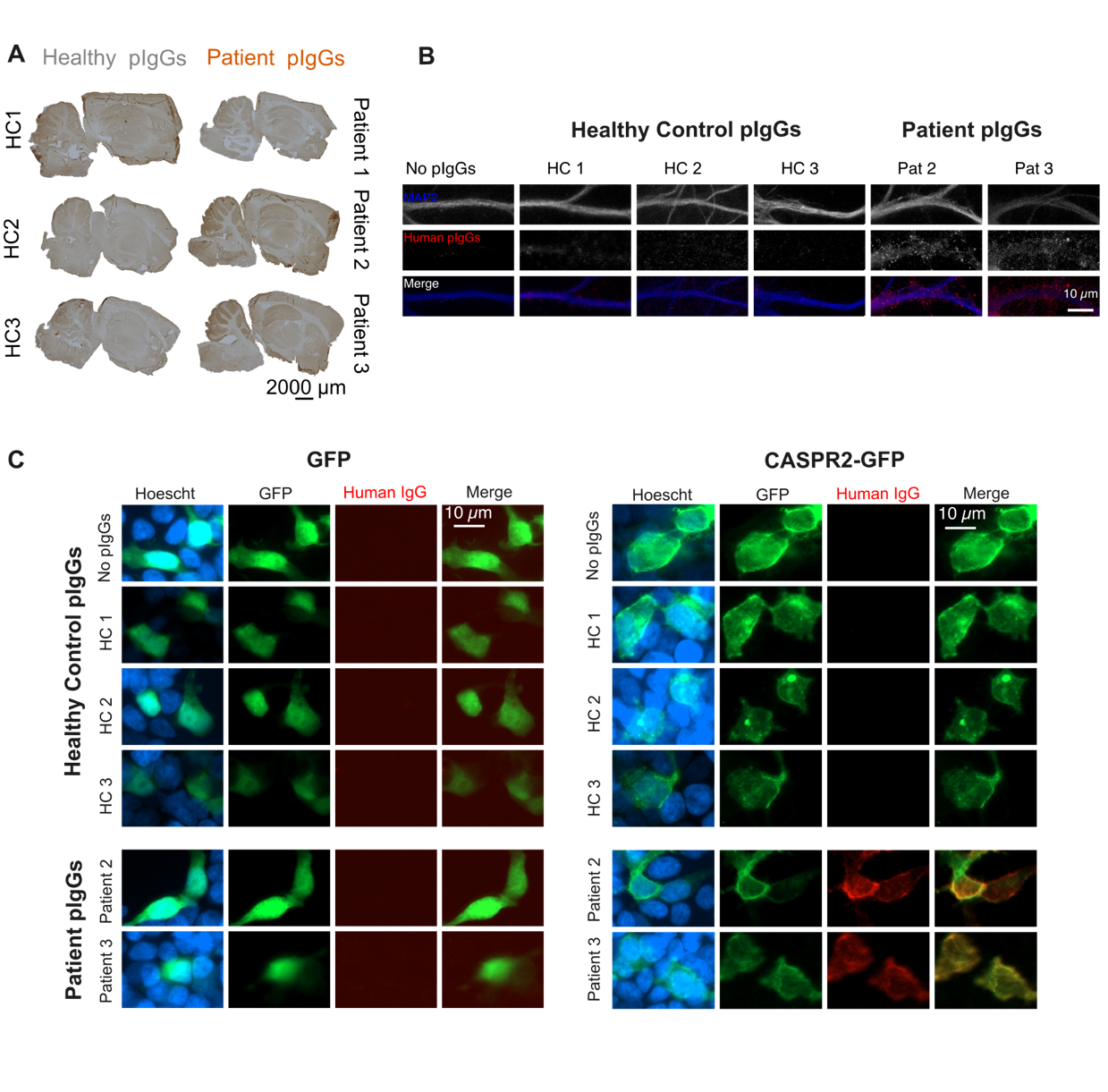
**Supplementary Figures**

**Fig. S1.** **CASPR2 autoantibodies found in the plasma of patients with neurological disease recognize CASPR2 at the cell surface.** Human samples containing healthy control IgGs and high-titer CASPR2 IgGs (two or three individuals per group) were incubated with brain tissue and cellular preparations to assess their surface binding and distribution. (**A**) Healthy control or Patient pIgGs samples were incubated for 1h at 1:100 with rat brain sagittal sections, followed by immunohistochemistry. Sections incubated with Patient plasma showed the characteristic binding pattern of CASPR2 Abs, whereas healthy control plasma did not reveal binding to brain sections. (**B**) Total fractions of human IgGs purified from the plasma of healthy control individuals or patients positive for CASPR2 Abs were incubated for 2h with DIV16 cultured hippocampal neurons. Neurons were fixed and stained for surface human IgG under non-permeabilizing conditions., then permeabilized and stained for the somatodendritic marker MAP2. Immunofluorescence images confirmed that Patient pIgGs bound to the neuronal surface, whereas healthy control pIgGs did not bind to neurons. (**C**) Total fractions of human IgGs purified from the plasma of healthy control individuals or patients positive for CASPR2 Abs were incubated for 2h iwith HEK cells expressing CASPR2-GFP or GFP. HEK cells were fixed and stained for surface human IgG under non-permeabilizing conditions. Immunofluorescence images confirmed that Patient pIgGs specifically recognized CASPR2, whereas healthy control pIgGs did not. *Abbreviations:* pIgGs, purified Immunoglobulins G; HC, healthy control.

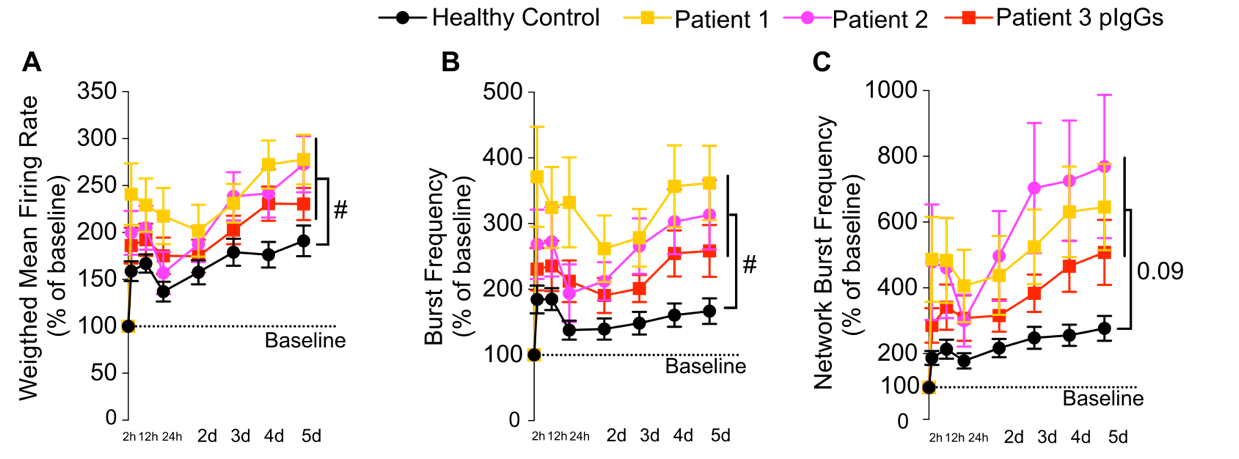

**Fig. S2. Patient-derived CASPR2 antibodies with neurological disease induce neuronal network hyperexcitability in rat cortical neurons.** Related to Fig. 1. Same experiments as in Fig. 1, with results shown for individual samples. (**A-C**) Patient pIgGs significantly increased (**A**) Weighted Mean Firing Rate, (**B**) Burst Frequency, and (**C**) Network Burst Frequency compared to Healthy pIgGs over 5 days of incubation with the human pIgGs. For all panels, statistical analysis was performed on the data shown in Table S6, using a two-way repeated measures ANOVA; # p < 0.05 indicates a significant main effect of pIgGs treatment. Results in all panels are presented as mean ± S.E.M. *Abbreviations:* pIgGs, purified Immunoglobulin G.

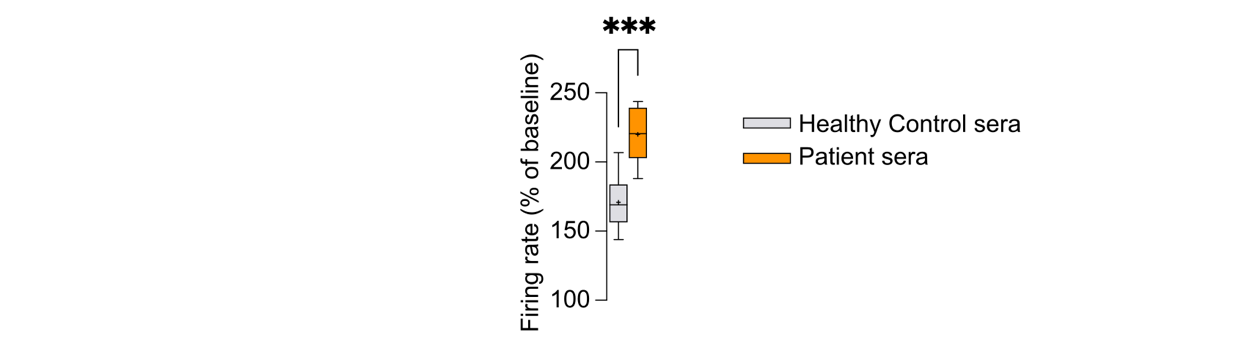

**Fig. S3. Sera containing CASPR2 antibodies with neurological disease induce neuronal increases the firing rate of iPSC-derived excitatory human neurons.** Related to Fig. 1. Human sera from patients or from control individuals was incubated at a dilution of 1:100 in iPSC-derived human neurons following monitoring of neuronal activity using MEA. Patient sera significantly increased the firing rate of iPSC-derived human neurons compared to healthy control sera over 6h of incubation, in MEA experiments. For healthy control sera, n= 6 wells, and for Patient sera, n = 12 wells. Statistical differences were determined using a unpaired t-test; ***p < 0.001 in comparison with healthy control sera. Results are presented as boxes showing the 25^th^ and 75^th^ percentiles and whiskers showing the range (minimum to maximum values corresponding to individual cells). The horizontal line within the boxes indicate the median. Crosses indicate the mean.

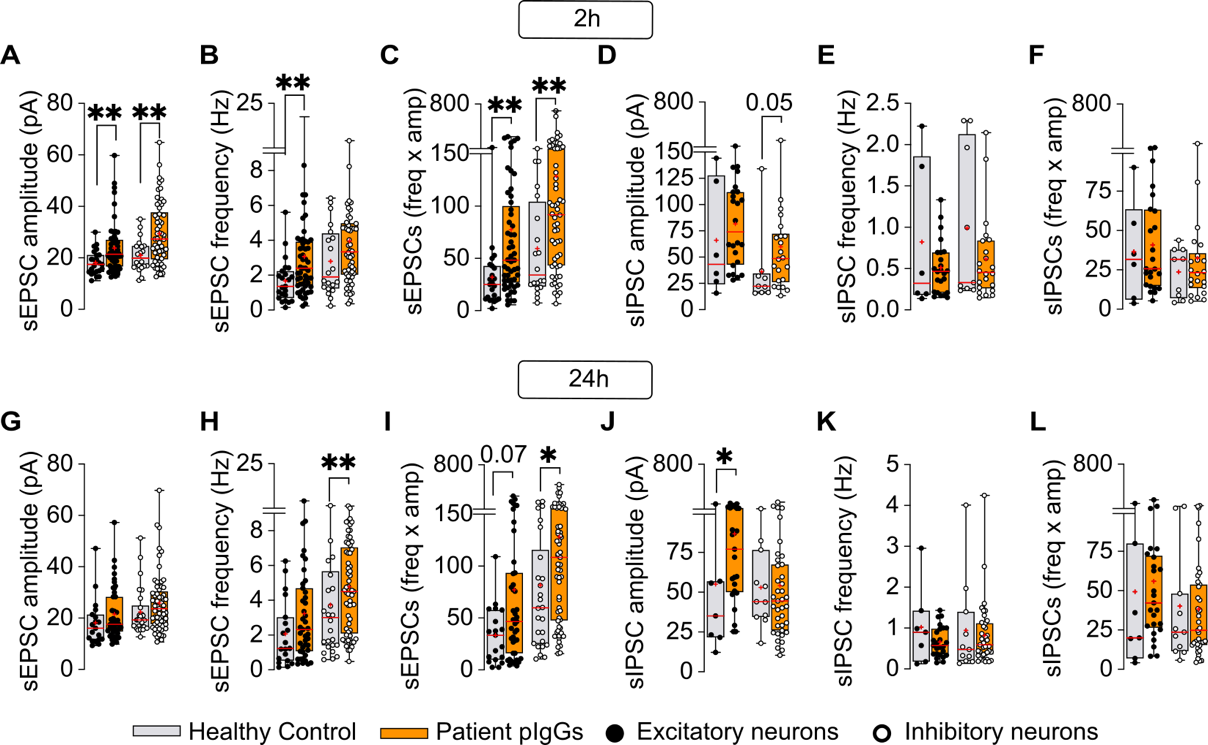

**Fig. S4. Patient-derived CASPR2 antibodies increase the amplitude and frequency of sEPSCs and the amplitude of sIPSCs in rat cortical neurons.** Related to Fig. 2. Same experiments as in Fig. 2, with results shown separately for inhibitory and excitatory neurons. (**A-C**) Patient pIgGs increased the sEPSCs amplitude (**A**) in excitatory and inhibitory neurons, the sEPSCs frequency (**B**) in excitatory neurons and the Amp x Freq (**C**) in both population of neurons after 2h and of incubation. (**D-E**) Patient pIgGs did not alter sIPSC amplitude (**D**), frequency (**E**), or Amp × Freq in either neuronal population after 2h of incubation. (**G-I**) Patient pIgGs did not alter the sEPSCs amplitude (**G**) in either neuronal population, increased the sEPSCs frequency (**H**) in inhibitory neurons and increased the Amp x Freq (**I**) in inhibitory neurons after 24h of pIgGs incubation. (**J-L**) Patient pIgGs increased the sIPSC amplitude (**J**) in excitatory neurons and altered neither the frequency (K), nor the Amp × Freq (**L**) in either neuronal population after 24h of incubation. For all panels, statistical analysis was performed on the data shown in Tables S7 and S8. Comparisons between conditions at 2h or 24h were performed using Mann–Whitney tests; * p *<* 0.05, ** p *<* 0.01, **** p *<* 0.0001 indicate significance in comparison to Healthy Control pIgGs. Where not stated, statistical significance was not found. Results are presented as boxes showing the 25^th^ and 75^th^ percentiles and whiskers showing the range (minimum to maximum values corresponding to individual cells). The horizontal line within the boxes indicate the median. Crosses indicate the mean. *Abbreviations:* pIgGs, purified Immunoglobulin G; Freq, frequency; Amp, amplitude; sEPSCs, spontaneous excitatory postsynaptic currents; sIPSCs, spontaneous inhibitory postsynaptic currents.

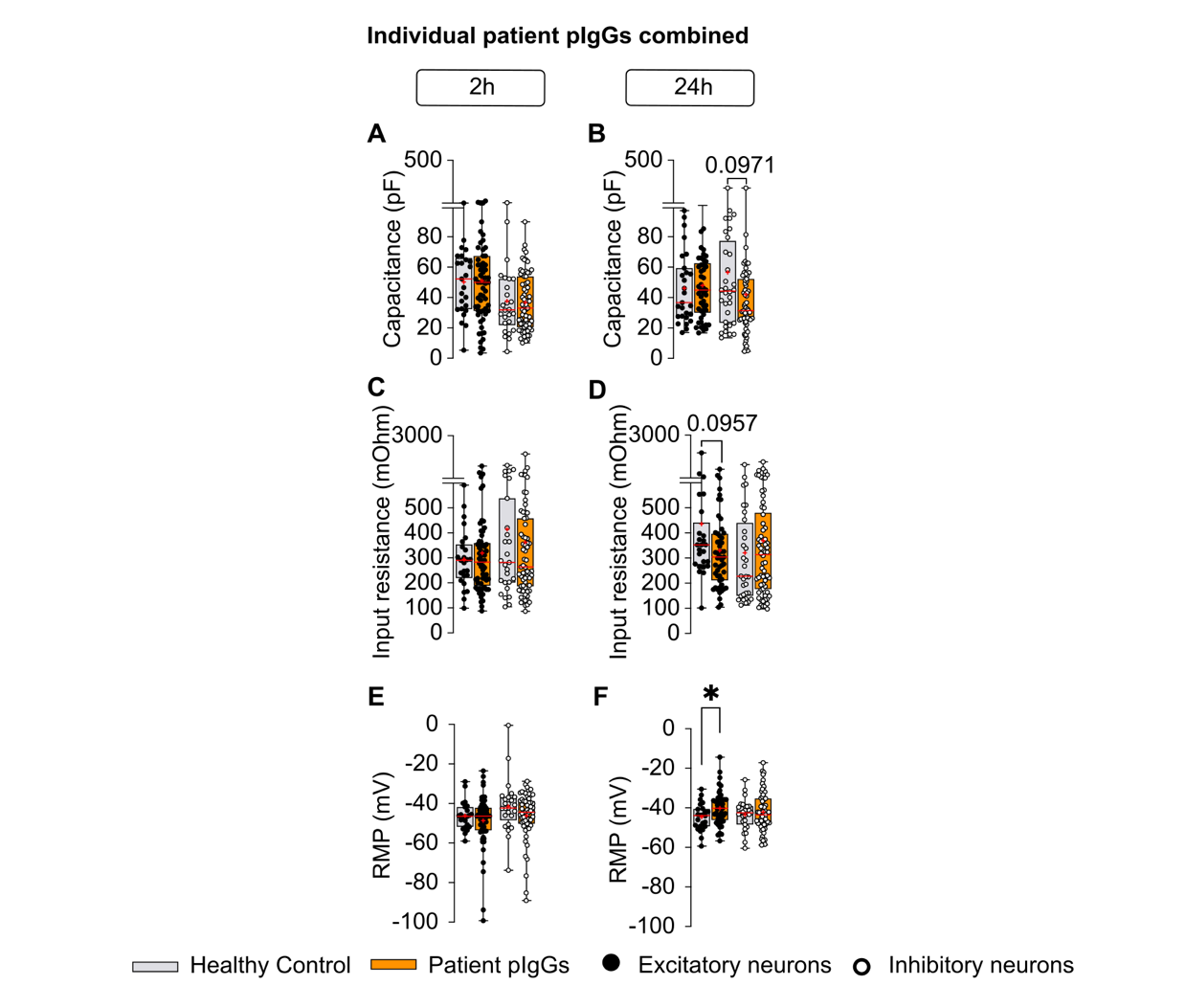

**Fig. S5. Patient-derived CASPR2 antibodies increase the resting membrane potential of excitatory neurons at 24h of human IgG incubation without altering other passive properties of neurons.** Related to Fig. 3. Same experiments as in Fig. 3a-c. Results shown separately for excitatory (black circles) and inhibitory (white circles) neurons following 2h (A-C) or 24h (D-F) of incubation. (**A,B**) Capacitance; (**C,D**) Input resistance and (**E,F**) RMP. Where not stated, statistical significance was not found. For all panels, statistical analyses were performed on data shown in Table S9 to compare between conditions within the same neuronal population. Results are presented as boxes showing the 25^th^ and 75^th^ percentiles and whiskers showing the range (minimum to maximum values corresponding to individual cells). The horizontal line within the boxes represents the median. Crosses indicate the mean. *Abbreviations:* pIgGs, purified Immunoglobulin G; RMP, Resting Membrane Potential.

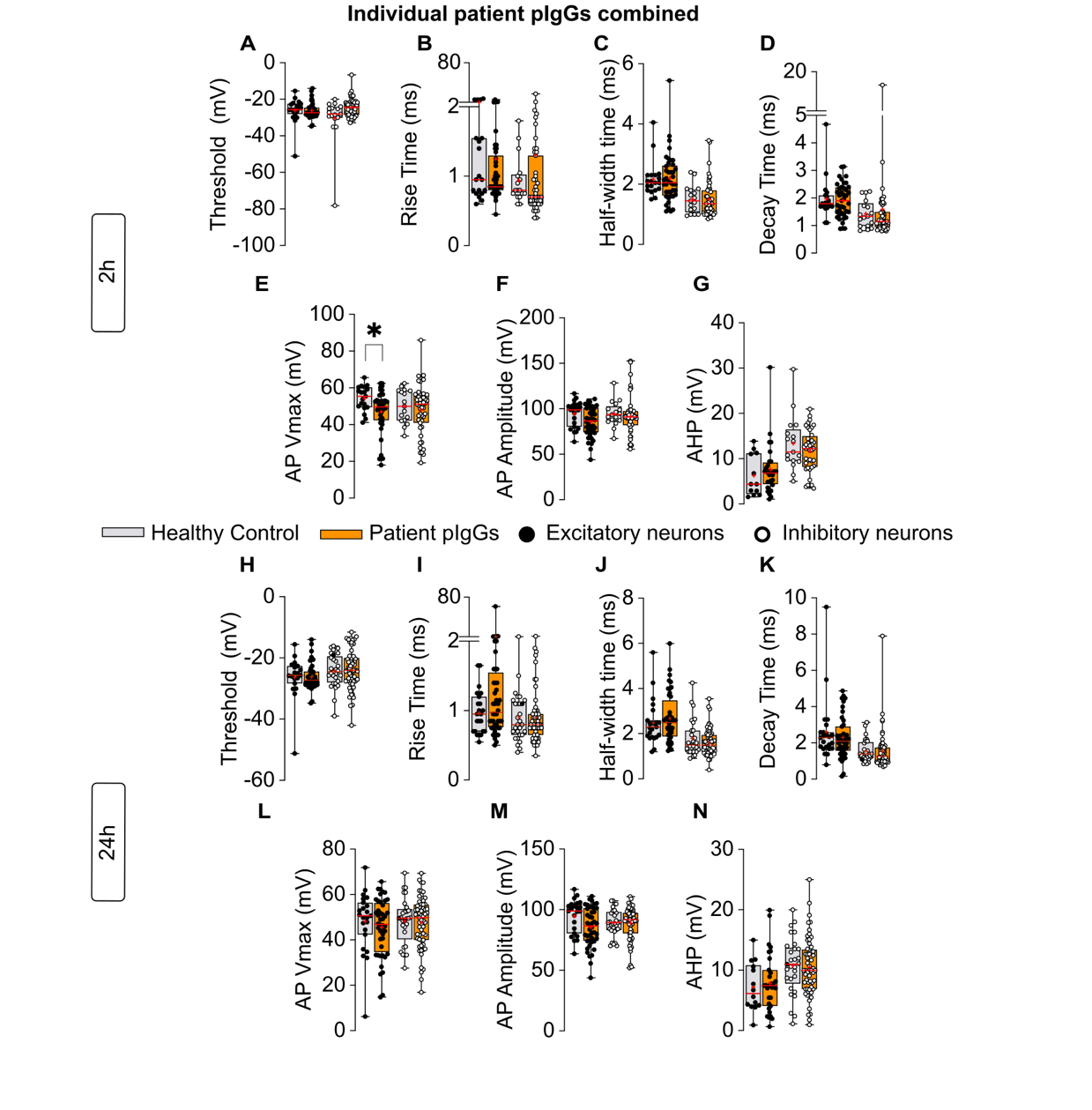

**Fig. S6. Patient-derived CASPR2 antibodies induce subtle changes in the action potential kinetics in inhibitory and excitatory cortical neurons.** (**A-N**) AP kinetics was analyzed using multiple parameters, including threshold (**A,N**), rise time (**B,I**), half-width (**C,J**), decay time (**D,K**), AP Vmax (**E,L**), AP amplitude (**F,M**), after-hyperpolarization (AHP) (**G,N**). Statistical analyses were performed on the data in the Table S10 between conditions at 2h or 24h within each neuronal population using the Mann-Whitney test; * p *<* 0.05. For all panels, where not stated, statistical significance was not found. Results are presented as boxes showing the 25^th^ and 75^th^ percentiles and whiskers showing the range (minimum to maximum values corresponding to individual cells). The horizontal line within the boxes indicates the median. Crosses indicate the mean. *Abbreviations:* pIgGs, purified Immunoglobulin G; HC, healthy control; AP, Action Potential; Vmax, maximum voltage; AHP, afterhyperpolarization peak.

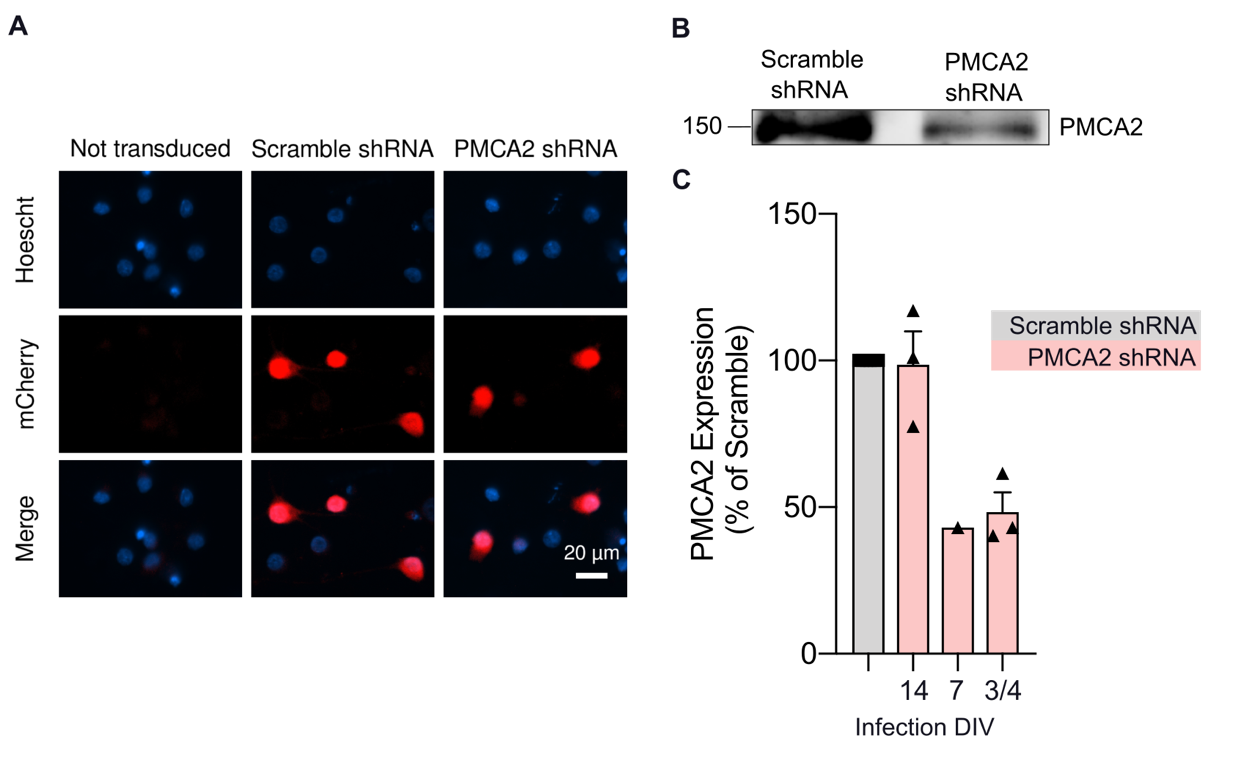

**Fig. S7. Validation of shRNA to knockdown** **PMCA2****.** (**A**) Images of rat cortical neurons transduced at DIV3-4 with lentiviruses enconding shRNAs to knockdown (KD) PMCA2 or a non-targeting (scramble) shRNA. mCherry^+^ neurons were efficiently transduced. (**B**) Representative WB showing that the PMCA2 shRNA efficiently decreased PMCA2 expression in rat cortical neurons when these were transduced at DIV3-4. (**C**) Levels of PMCA2 expression after KD in rat cortical neurons, when transduced at different DIVs.

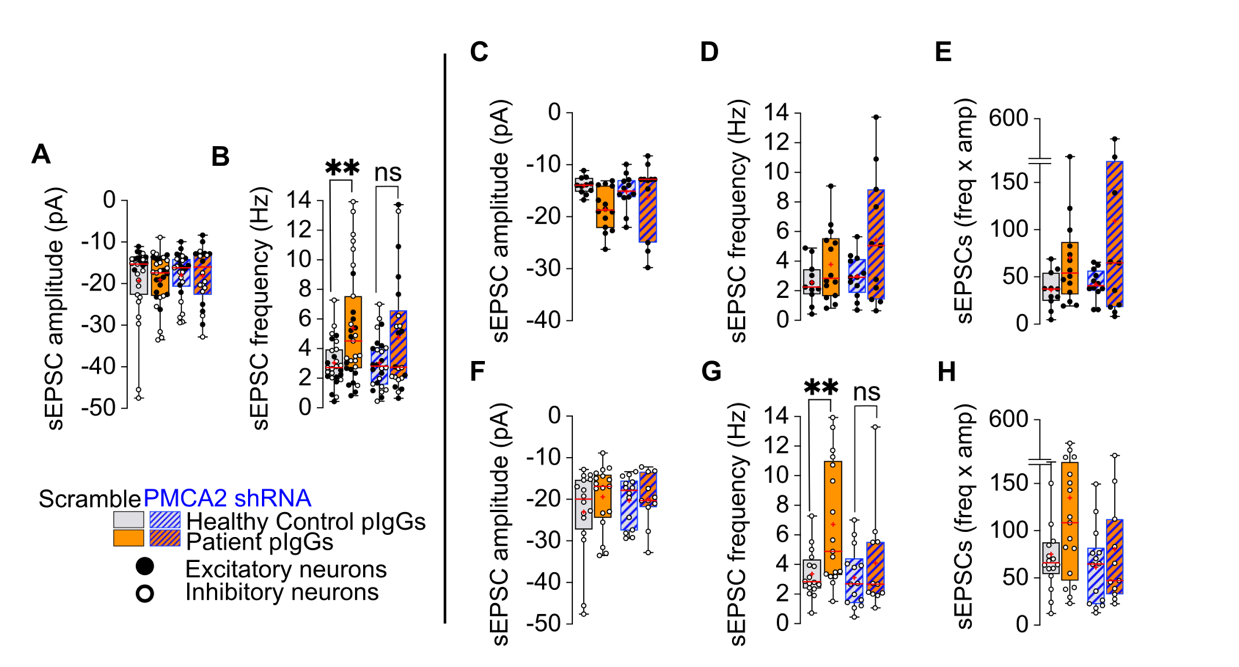

**Fig. S8. PMCA2 silencing prevents patient CASPR2 antibodies-induced increase in the frequency of sEPSCs in cortical neurons.** Related to Fig. 7, with results shown separately for each metric. (**A,B**) PMCA2 KD prevented the increase in sEPSC amplitude (**A**) and frequency (**B**) following 24h of incubation with the human pIgGs when considering both excitatory and inhibitory neurons. Statistical comparisons were performed on sEPSCs data [N = 5 independent experiments: n(Scramble HC pIgGs) = 29 neurons, n(Scramble Patient pIgGs) = 31 neurons, n(PMCA2 shRNA HC pIgGs) = 26 neurons, n(PMCA2 shRNA Patient pIgGs) = 22 neurons]. (**C-E**) Patient pIgGs did not affect the amplitude (**C**), the frequency (**D**) nor the frequency x amplitude (**E**) of sEPSCs following 24h of incubation in excitatory neurons. (**F-H**) Patient pIgGs did not affect the amplitude (**F**), the frequency (**G**), nor the frequency x amplitude (**H**) of sEPSCs following 24h of incubation in inhibitory neurons. Statistical comparisons were performed on sEPSCs data [N = 5 independent experiments: n(Scramble HC pIgGs) = 9 neurons, n(Scramble Patient pIgGs) = 12 neurons, n(PMCA2 shRNA HC pIgGs) = 9 neurons, n(PMCA2 shRNA Patient pIgGs) = 4 neurons] using a two-way ANOVA to compare differences between conditions; * p *<* 0.05 indicates significance compared to Healthy Control pIgGs. Where not stated, statistical significance was not found. Results are presented as boxes showing the 25^th^ and 75^th^ percentiles and whiskers showing the range (minimum to maximum values corresponding to individual cells). The horizontal line within the boxes represents the median. Crosses indicate the mean. *Abbreviations:* pIgGs, purified Immunoglobulin G; Freq, Frequency; Amp, Amplitude; sEPSCs, spontaneous Excitatory Postsynaptic Currents.

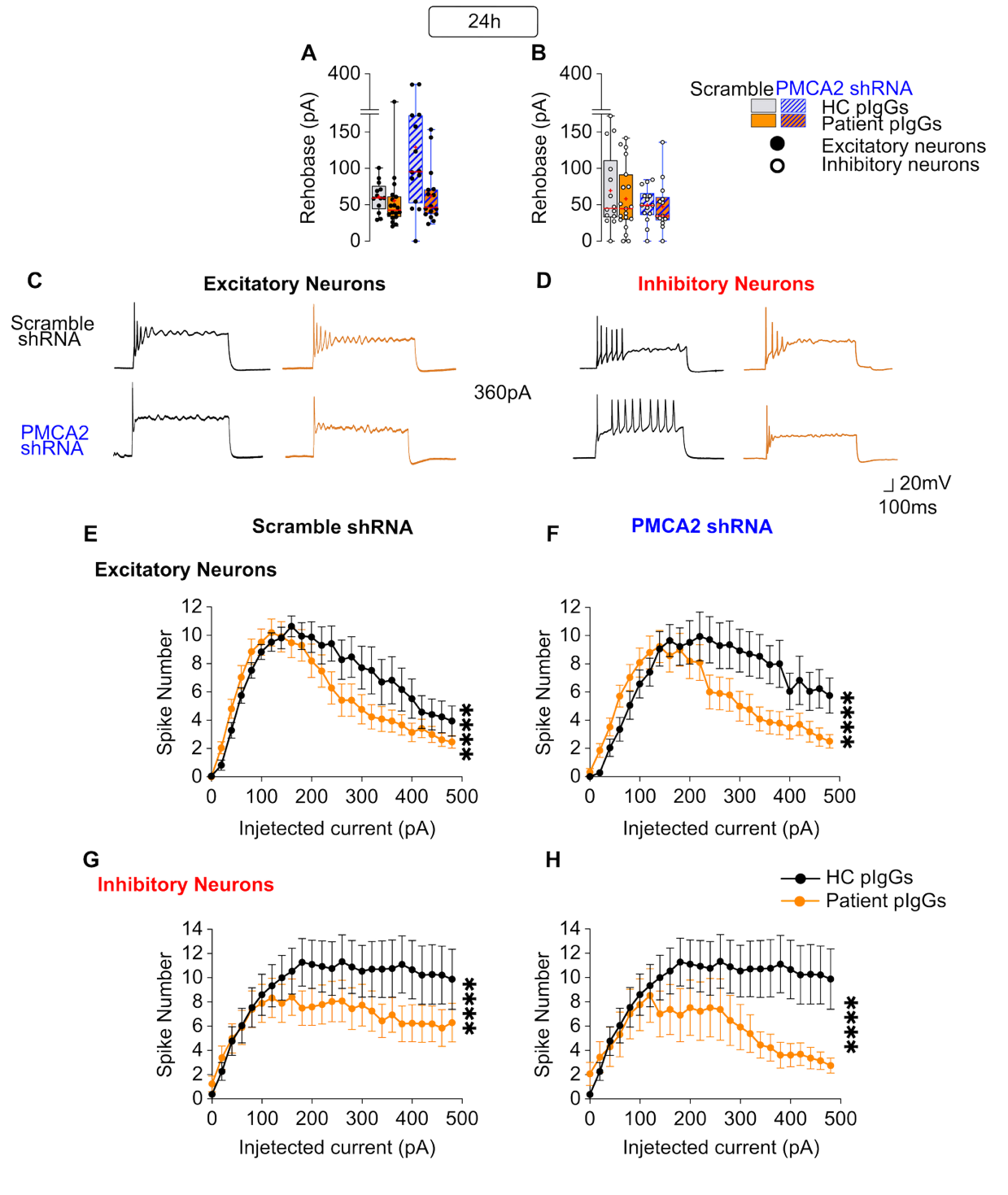

**Fig. S9. Silencing PMCA2 does not rescue the excitability changes induced by the patient-derived CASPR2 antibodies on inhibitory and excitatory neurons at 24h.** DIV14-15 neurons were incubated for 24h with pIgGs from healthy controls or individual patient CASPR2 pIgGs at 20µg/mL, prior to whole-cell patch-clamp recordings. Results are shown separately for excitatory (black circles) and inhibitory (white circles) neurons. (**A,B**) The rheobase in excitatory (**A**) and inhibitory (**B**) neurons was not affected by PMCA2 KD after incubation with patient pIgGs . Statistical comparisons were performed on excitatory neurons [N = 5 independent experiments: n(Scramble HC pIgGs) = 13 neurons, n(Scramble Patient pIgGs) = 18 neurons, n(PMCA2 shRNA HC pIgGs) = 13 neurons, n(PMCA2 shRNA Patient pIgGs) = 16 neurons]. Statistical comparisons were performed on inhibitory neurons [N = 5 independent experiments: n(Scramble HC pIgGs) = 13 neurons, n(Scramble Patient pIgGs) = 17 neurons, n(PMCA2 shRNA HC pIgGs) = 13 neurons, n(PMCA2 shRNA Patient pIgGs) = 13 neurons] using a two-way ANOVA followed by Tukey’s multiple-comparisons post hoc test. (**C-F**) Patient pIgGs decreased the excitability of excitatory and inhibitory neurons in Scramble (**C,E**) and PMCA2 (**D,F**) shRNA conditions. Statistical comparisons performed on current clamp data in excitatory neurons [N = 5 independent experiments: n(Scramble Healthy Control pIgGs) = 17 neurons, n(Scramble Patient (pIgGs) = 21 neurons; n(PMCA2 shRNA Healthy Control pIgGs) = 17 neurons, n(PMCA2 shRNA Patient pIgGs) = 21 neurons] and on inhibitory neurons [N = 5 independent experiments: n(Scramble Healthy Control pIgGs) = 18 neurons, n(Scramble Patient pIgGs) = 20 neurons, n(PMCA2 shRNA Healthy Control pIgGs) = 18 neurons, n(PMCA2 shRNA Healthy Control pIgGs) = 13 neurons] using a non-parametric two-way ANOVA Test. **** p *<* 0.0001 for main differences between groups. Where not stated, statistical significance was not found. Results are presented as boxes showing the 25^th^ and 75^th^ percentiles andwhiskers showing the range (minimum to maximum values corresponding to individual cells). The horizontal line within the boxes represents the median. Crosses indicate the mean. *Abbreviations:* HC, healthy control; pIgGs, purified Immunoglobulin G.

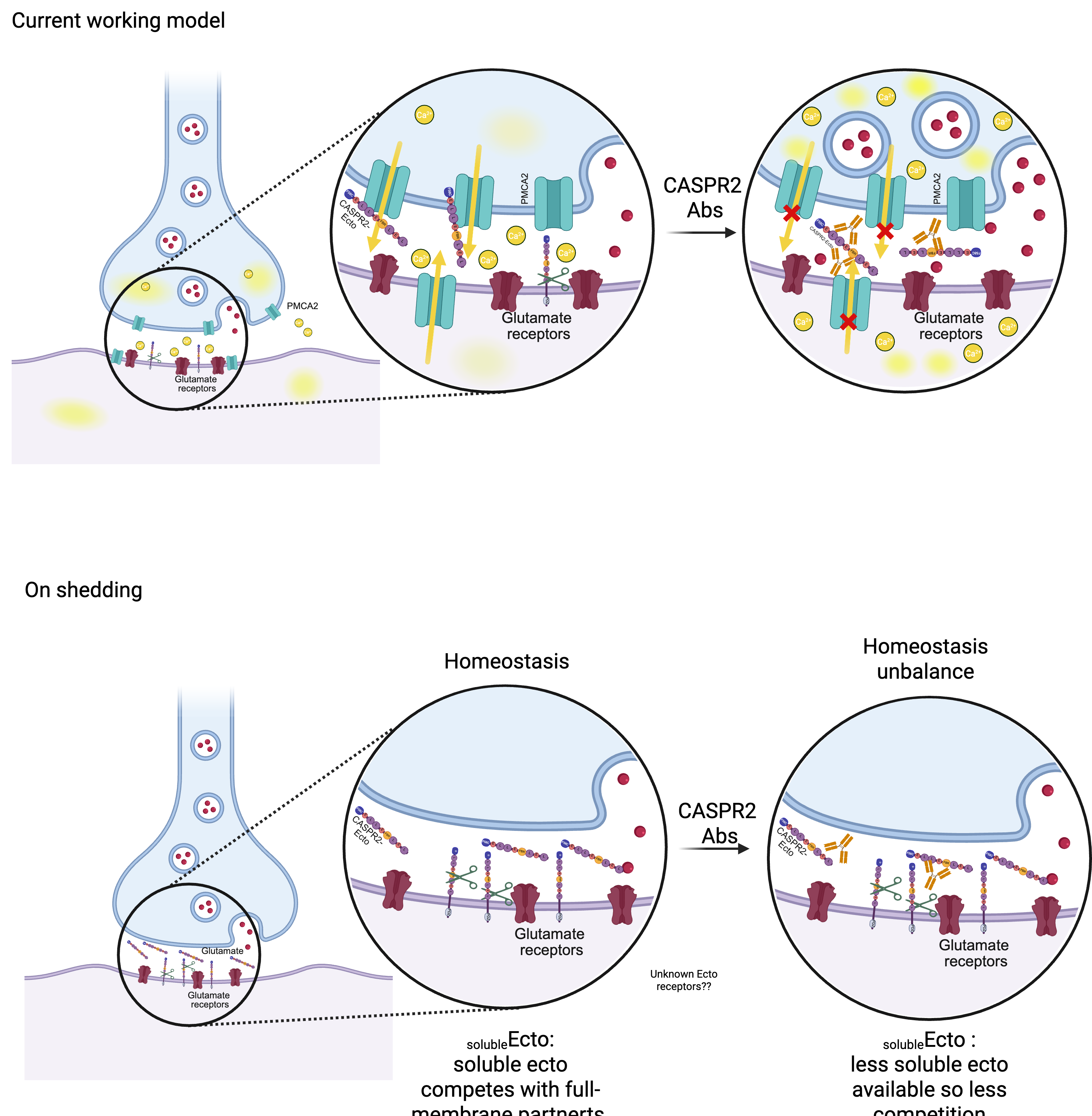

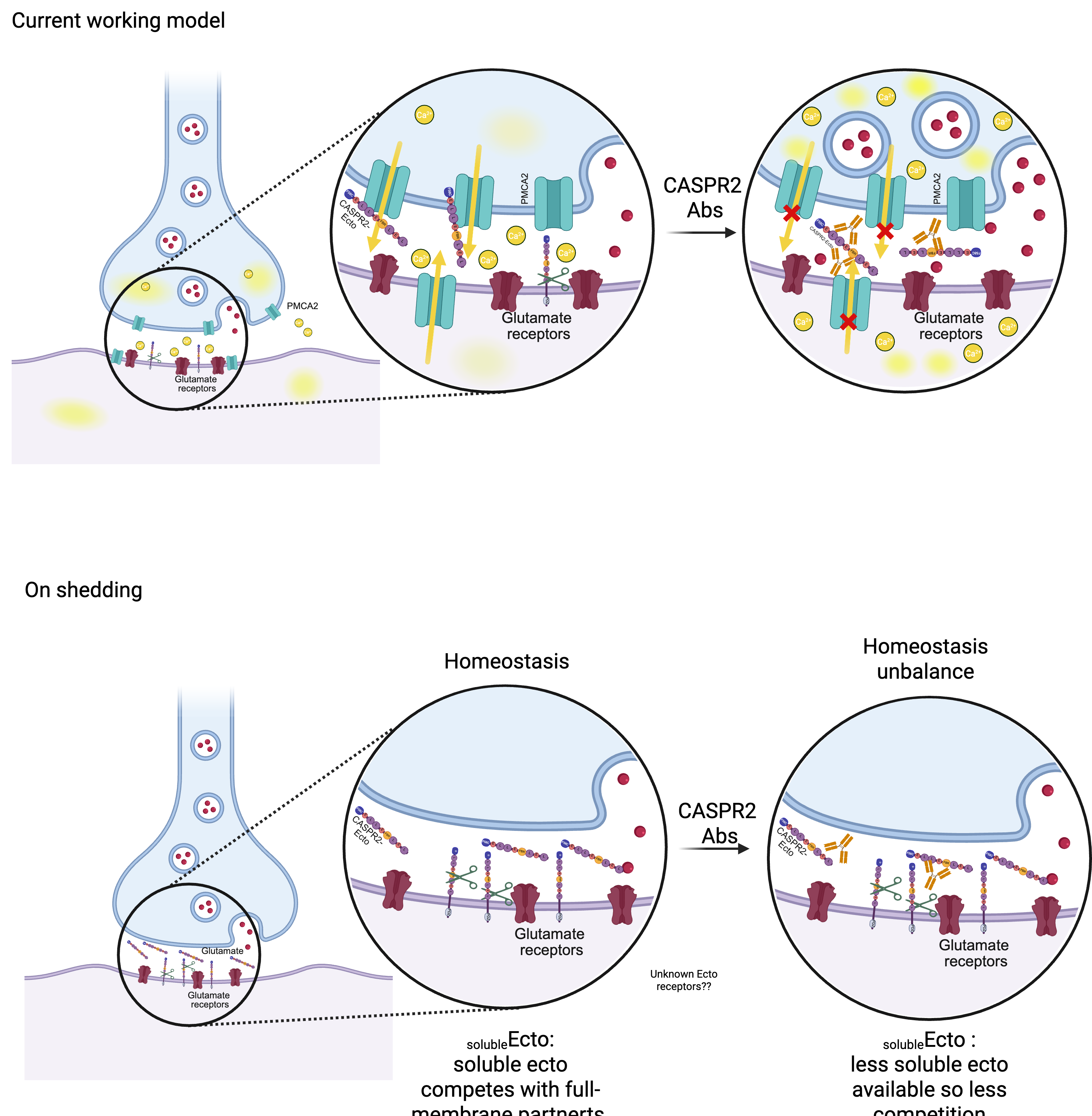

CASPR2 ectodomain

PMCA2

PMCA2

**Fig. S10. Model of CASPR2 autoantibody pathogenesis through sequestration of the shed CASPR2 ectodomain.** Our data support a model in which disruption of the shed CASPR2 ectodomain contributes to the hyperexcitability induced by CASPR2 antibodies. Under physiological conditions, the shed ectodomain constitutively binds to PMCA2, maintaining calcium levels and neuronal activity within a physiological range. In patients with CASPR2 autoimmunity, autoantibodies bind and sequester the soluble CASPR2 ectodomain, reducing the pool of free ectodomain available to interact with PMCA2. This loss of interaction lowers PMCA2 activity, impairing calcium clearance. The resulting rise in intracellular calcium increases neuronal network activity, leading to hyperexcitability.

**Supplementary Materials and Methods**

**Rat primary neuronal cultures**

Briefly, the brains were dissected in Ca^2+^- and Mg^2+^-free Hank’s balanced solution (HBSS) containing (in mM): 5.36 KCl, 0.44 KH_2_PO_4_, 137 NaCl, 4.16 NaHCO_3_, 0.34 Na_2_HPO_4_.2H_2_O, 5 glucose, 1 pyruvic acid, 10 4-(2-hydroxyethyl)-1-piperazineethanesulfonic acid (HEPES) and 0.001% (w/v) phenol red. Brain tissues were enzymatically digested with 0.2% (w/v) trypsin in HBSS, supplemented with deoxyribonuclease (0.01% v/v; Sigma-Aldrich, USA: #DN25), for 15 min in a water bath at 37ºC. After incubation, the supernatant was discarded and the trypsin action was stopped with serum-containing plating medium composed of minimum essential medium (MEM; Sigma Aldrich, USA: #M0268) supplemented with 10% (v/v) heat-inactivated horse serum, 0.22% (w/v) NaHCO_3_, 0.6% (w/v), and 1 mM pyruvic acid. Next, the tissue was mechanically dissociated, and cell density was determined with tryptan blue.

High-density cultures (90 × 10³ cells/cm²) for biochemical studies were prepared in 6-well plates. Semi-dense cultures (53 × 10³ cells/cm²) used for electrophysiological and calcium-imaging experiments were prepared on glass coverslips in 12-well plates. Cells were left to adhere for 2-4 hours in a humidified incubator of 5% CO_2_/95% air at 37°C. Afterwards, the medium was replaced with Neurobasal culture medium (NBM; GIBCO Invitrogen, USA: #21103049) supplemented with SM1 (Stem Cell Technologies, Canada: #05711), Glutamax (GIBCO Invitrogen, USA: #35050-061), and 0.012% (w/v) gentamicin (GIBCO Invitrogen, USA: #15750060). Cultures were fed weekly by replacing one third of the medium volume with fresh one without glutamate. Low-density Banker cultures for LCBA experiments were prepared as previous published works (*48*). Cells were plated on glass coverslips at a density of 9500 cells/cm^2^. After 2-4 hours, the coverslips were flipped over an astroglial feeder layer in Petri dishes or 12-well plates containing NBM and fed weekly. For Microelectrode Arrays (MEA) recordings, the cell suspension was supplemented with laminin (Sigma Aldrich, USA: #L2020) at a final concentration of 10 *µ*g/mL. A drop of 10 *µ*l of this mixture was then plated on top of the electrode array in a 24-well MEA plate (Axion Byosystems, USA: #M384-tMEA-24W) previously coated with. Cells were plated at a density of 5.3 x 10^3^ cells/mm^2^ (corresponding to 30 x 10^3^ cells plated per well) or at a density of 13.96 x 10^3^ cells/mm^2^ (corresponding to 80 x 10^3^ cells per well) and NBM was added to each well after allowing cells to adhere. MEA cultures were fed weekly starting at day in vitro (DIV) 7 by replacing one third of the conditioned medium with fresh prewarmed BrainPhys (STEMCELL Technologies, USA: # 05790) supplemented with SM1 (Stem Cell Technologies, Canada: #05711) and 0.012% (w/v) Gentamicin (GIBCO Invitrogen, USA: #15750060) and then at DIV14 with NBM. For hippocampal cultures, the NBM was also supplemented with 25 *µ*M glutamate on the isolation day. To prevent overgrowth of the glial layer, 10 *µ*M 5-Fluoro-2-_´_deoxyuridine (FDU) (Sigma-Aldrich, USA: #50-91-9) was added to all cultures after 3 or 4 DIV, except for the high-density cultures. Cultures were maintained in a humidified incubator of 5% CO_2_ 95% air at 37°C.

**Human Embryonic Kidney (HEK293T) cell cultures**

HEK293T cells (ATCC: #CRL-3216) were grown in Dulbecco’s Modified Eagle Medium (DMEM) (Sigma Aldrich, USA: #D5648) supplemented with 44 mM NaHCO_3_, 10% (v/v) of heat-inactivated fetal bovine serum (FBS) and 1% (v/v) penicillin/streptomycin (PenStrep) (GIBCO Invitrogen, USA: #15140122) at pH 7.2. Cells were maintained in a humidified incubator containing 5% CO_2_/ 95% air at 37°C. Cells were split every 3-4 days using 0.25% (w/v) Trypsin-EDTA (GIBCO Invitrogen, USA: #25200056) and re-plated in fresh medium at a 1:20 dilution. Cells were plated at different densities depending on the specific experiment to be performed (see supplementary material for details).

For live cell-based assays (LCBAs), HEK293T cells were seeded at a density of 160 × 10³ cells/well on 18 mm poly-D-lisine (PDL)-coated glass coverslips in 12-well plates, reaching 60% confluency by the day of transfection. For low-titer AAV production, cells were seeded at a density of 300 × 10³ cells/well on PDL-coated 6-well plates, reaching 80% confluency at transfection. For high-titer lentivirus production, cells were seeded on 100 mm petri dishes and transfected at 70% confluency.

**hiPSC differentiation into NPCs and neurons**

To start EB formation*,* day 0, AggreWell™800 24-well (Stemcell Technologies) plates were pretreated with Anti-Adherence Rinsing Solution (Stemcell Technologies) and centrifuged to clear bubbles from the microwells. Each well was then filled with 1 mL of warm induction medium containing 10 μM Y-27632. hiPSCs were dissociated into single cells with Gentle Cell Dissociation Reagent and counted with trypan blue (Sigma-Aldrich). Cells were pelleted and resuspended at 3 x 10⁶ cells/mL in induction medium with 10 μM Y-27632. One milliliter of suspension (3 x 10⁶ cells) was added per well, targeting 10,000 cells per microwell. Plates were centrifuged to pellet the cells. A daily half medium change was done with fresh induction medium including 10 μM Y-27632. On day 8, the EBs were harvested with wide-bore tips and passed through a 37 μm reversible strainer. They were replated intact onto Matrigel coated 6-well plates. Medium was fully replaced each day through day 11. On Day 12, rosette selection was carried out by first washing with DMEM/F-12 and then incubating cells with STEMdiff™ Neural Rosette Selection Reagent (Stemcell Technologies) for 1.5 h at 37°C. Rosettes were detached by gently washing with DMEM/F-12 and pelleted. They were resuspended in induction medium and replated onto freshly coated 6-well plates. Medium was changed daily through days 17 to 19. Between day 17 and 19 the NPCs were passaged with ACCUTASE™ (Stemcell Technologies). They were replated in STEMdiff™ Neural Progenitor Medium with daily medium changes. Later passages were done after 6 to 7 days. NPCs were cryopreserved in STEMdiff™ Neural Progenitor Freezing Medium (Stemcell Technologies). Forebrain-type neurons were generated from hiPSC-derived NPCs with the STEMdiff™ Forebrain Neuron Differentiation Kit and the STEMdiff™ Forebrain Neuron Maturation Kit (Stemcell Technologies).

**Transfection of HEK293T cells**

To produce adeno-associated viruses (AAVs), HEK293T cells were transfected with a pDeltaF6 helper vector, a vector encoding for Rep/Cap genes and a vector expressing either hDXL-linker-mCherry, mDLX-GFP or GCaMP6f using a DNA:PEI ratio of 1:4 (µg:µL) was used.Prior to transfection, the HEK293T cells medium was replaced with DMEM supplemented with 2% FBS and 1% PenStrep. Finally, the DNA mix was added to the cells, and cells were left for 3 days in an humidified atmosphere with 5% CO_2_ at 37ºC. To produce lentiviruses (LVs), the following constructs were used: a vesicular stomatitis virus glycoprotein (VSVG) construct encoding an envelope protein, a PAX2 packaging construct and the constructs to express scramble (control) or PMCA2 shRNAs. To prepare the mixes, 100 *µ*g of each construct were diluted in prewarmed Opti-MEM (Thermo Scientific, USA: #22600134) supplemented with 29 mM NaHCO_3_ and 1% PenStrep and added PEI to a final DNA:PEI ratio of 1:6 (µg:µL). Cells were allowed to produce the viruses for 3 days. For the LCBAs, to prepare the DNA precipitates, the DNA was diluted in ddH2O and a 2M CaCl_2_ solution was then added to a final concentration of 2 mM. The mix was vortexed and added dropwise to HEPES buffered saline solution (HEBS in mM: 50 HEPES, 280 NaCl, 1.5 Na2HPO4, pH 7.05). The DNA precipitates were incubated for 5h with the cells before the medium was replaced with fresh DMEM culture medium supplemented with 1% PenStrep and 10% FBS and cells were returned to the same incubating conditions for approximately 48h, at which point the LCBAs were performed.

**Purification of viruses**

To purify low-titer adeno-associated viruses (AAVs), HEK293T medium was first filtered to remove cell debris. Afterwards, the medium was concentrated using 100 kDa MW cut-off Amicon® Ultra filter columns (Millipore, USA: #UFC5100). The columns were equilibrated in phosphate-buffered saline (PBS) supplemented with Magnesium and Potassium (PBS-MK) buffer, followed by centrifugation for 2 min at 14000xg. Subsequently, the conditioned medium was loaded in a column followed by centrifugation for 3 min at 14000xg. This step was repeated as many times as needed to allow all the media to flow through the column. The column was then washed with PBS-MK, followed by a 3 min centrifugation. This step was repeated three times. To prevent aggregation of viral particles, the column was loaded with 0.01% pluronic in PBS to a final volume of 50 µL and centrifuged. To collect the virus concentrated in the column, the column was inverted in a clean collection tube and spined down for 1 minute. The viruses were then aliquoted and stored at -80°C until future use. We validated the viral infections by confirming colocalization of the mCherry signal (from the hDLX construct) with GABA immunoreactivity in neurons (data not shown).

To purify high-titer lentivirus (LVs), HEK293T conditioned medium was centrifuged for 10 min at 2000 rpm and at 4°C and filtered (VWR, USA: #514-0329) to remove cell debris. To concentrate the viruses, the medium was ultracentrifuged in a sucrose gradient. Briefly, an ultracentrifuge tube (Beckman Coulter, USA: #082326823) was loaded with 20 mL of conditioned medium followed by careful pipetting of 4 mL of a 20% sucrose solution into the bottom of the tube, creating a visible layer and forming a sucrose gradient. The medium was ultracentrifuged for 2h at 20000 rpm at 4°C (Beckman Coulter, USA: OptimaTM L-100 XP ultracentrifuge, Rotor SW28). Afterwards, the supernatant was carefully aspirated and tubes were allowed to dry for 10 min. Viruses were then resuspended in PBS, aliquoted and stored at -80°C until future use.

PMCA2 LV titers were determined with a titration assay using rat cortical neurons. PMCA2 knockdown (KD) was validated by transducing cortical cells at DIV 3, 7, and 14 with the shRNA pair of viral constructs at a final moiety of infection (MOI) of 5 (representing about 80% neuronal infection). WB experiments carried out with a PMCA2 commercial antibody confirmed that PMCA2 expression was reduced to 50% in KD condition when transduction took place at DIV3-7 (Fig. S7).

**Purification of human IgG from patient plasma**

Protein G beads were washed twice with 3x bead volume of DPBS (ThermoFisher, USA: 14190250) prior to use. Plasma was incubated with Protein G beads at a 5:6 plasma-to-bead volume ratio to ensure complete IgG capture and incubated overnight at 4°C with agitation. The plasma-bead mixture was centrifuged at 1,000 × g for 10 min, and the supernatant was removed. The beads were transferred to Pierce Centrifuge Columns (ThermoFisher, USA: 89868), centrifuged at 1,000 × g for 1 min, and washed three times with 500 µL DPBS, centrifugation at 1000 × g for 1 min. Bound IgG was eluted three times by adding 300 µL Pierce IgG Elution Buffer (ThermoFisher, USA: 21009) to the beads, incubating for 1 min, and centrifuging at 1,000 × g for 2 min into separate tubes containing 30 µL 1 M Tris (pH 9.0) (ThermoFisher, USA: J62085.K2). Purified IgG was buffer-exchanged into DPBS and concentrated using a 30-kDa Amicon Ultra Centrifugal Filter (Millipore Sigma, USA: UFC803024). The final IgG concentration was determined by absorbance at 280 nm using a NanoDrop spectrophotometer.

**Immunocytochemistry general protocol**

Cells were fixed in 4% (v/v) paraformaldehyde (PFA)/ 4% (w/v) sucrose in PBS for 15 min at RT, followed by 6 PBS washes. Cells were permeabilised with 0.25% Triton in PBS for 5 min at 4°C, followed by 6 washes in PBS. To block nonspecific staining, cells on coverslips were incubated in 10% (w/v) bovine serum albumin (BSA) in PBS for 30 min at 37°C. To immunolabel the proteins of interest, neurons were incubated with a mix of primary antibodies diluted in 3% (w/v) BSA in PBS 2h at 37°C or overnight (ON) at 4°C. Afterwards, cells were washed with PBS and incubated with fluorophore-conjugated secondary antibodies for 1h at 37°C. Coverslips were washed in PBS and mounted with DAKO (Glostrup, Denmark: #S302380-2) mounting medium on glass microscope slides. To label extracellular epitopes, ICCs were performed in a similar manner using live-staining and under non-permeabilising conditions.

**CASPR2 Live Cell Binding Assays**

LCBAs were carried out to assess the surface binding of human IgGs to HEK293T cells expressing CASPR2. HEK293T cells expressing EGFP-tagged CASPR2 or EGFP (as control) were incubated with plasma or total purified IgGs from patients or healthy controls at a concentration of 20 µg/mL for 2h. Subsequently, cells were fixed and quickly washed in warm PBS. Cells were then blocked with xx% BSA in PBS, followed by incubation with anti-human IgG secondary antibody (in BSA 3% (w/v) in PBS) for 1h at 37°C. After incubation, the coverslips were washed 6 times in PBS and mounted with DAKO containing 1µg/mL of Hoechst.

**Human IgG binding to the surface of live neurons**

To test whether purified IgG samples from CASPR2-Abs^+^ patient or healthy controls bound to the neuronal surface, DIV16 primary neurons were incubated with patient or control purified IgGs at a IgG concentration of 20 µg/mL for 2h in a humidified atmosphere with 5% CO_2_ at 37°C. After incubation, unbound IgGs were washed out by quickly rinsing coverslips in warm PBS. Cells were fixed and blocked as described above, and then incubated with anti-human fluorophore-conjugated secondary antibody ON, at 4°C. On the following day, coverslips were washed in PBS and cells were permeabilized and re-blocked. To immunolabel the somatodendritic compartments, coverslips were incubated with anti-MAP2 primary antibody for 2h at 37°C, washed in PBS and then incubated with a proper secondary antibody for 45 min at 37°C. Finally, coverslips were washed and mounted in DAKO, in which the DNA dye Hoechst (1 µg/ µL) had been diluted.

**Human IgG binding to brain slices**

Briefly, 7 µm frozen sagittal sections obtained from Wistar rats were rehydrated with Trizma-buffered saline (TBS) for 5 min. Next, sections were incubated with 0.3% H_2_O_2_-TBS for 15 min at 4°C to quench residual peroxidase activity. After 3 washes with TBS, the slices were blocked with TBS-10% normal goat serum (NGS) for 1h. Brain sections were incubated with patient CASPR2 IgGs or controls at a 1:100 dilution and left ON at 4 °C. On the subsequent day, slices were washed again 3 times with TBS and incubated for 2h with a secondary biotinylated anti-human IgG 1:1000 in TBS-5% NGS at RT, washed again and then incubated with an Avidin-Biotin kit (ABC,VectorLabs, USA: #PK-6100). After 3 more washes, the chromogenic reaction was developed with a Vectastain 3,30-diamenobenzydine (DAB) kit (Enzo Life Sciences, USA: #NC1323559) for 6-7 min. Subsequently, the slides were dehydrated by increasing ethanol concentration solutions and xylene. Finally, the slices were mounted using DPX mounting medium and air-dried under a fume hood, ON.

**Preparation of protein extracts**

Whole cell lysates for Western Blot (WB) analysis were prepared from DIV15 or DIV21 cortical neurons. Briefly, cells were lysed in ice-cold radioimmunoprecipitation assay buffer (RIPA), containing 150 mM NaCl, 1.0% (v/v) Triton X-100, 0.5% (w/v) sodium deoxycholate, 0.1% sodium dodecyl sulphate (SDS) and 50 mM Tris, pH 8.0], supplemented with a mix of protease inhibitors [1 mM DTT, 0.2 mM phenylmethylsulfonyl fluoride (PMSF), 1 µg/ml Chymostatin, Leupeptin, Antipain, Pepstatin-CLAP] and phosphatase inhibitors (50 mM sodium fluoride, 1 mM sodium ortovanadate). Lysates were left in orbital rotation for 30 min at 4°C followed by a 15 min centrifugation at 21.100 x g. Protein levels were quantified using the BCA assay kit (Pierce, Thermo Fisher Scientific, USA: #23225). Protein extracts were denatured with a 5x concentrated Laemmli Sample buffer containing 62.5 mM Tris-HCl (pH 6.8), 10% (v/v) Glycerol, 2% (w/v) SDS, 0.01% (w/v) bromophenol blue, 5% (v/v) β-mercaptoethanol]. Extracts were left unboiled or were boiled for 5 min at 95°C and stored at -20°C until being used.

**Precipitation of proteins in the extracellular medium**

Endogenous soluble CASPR2 ectodomain in cultured neurons was detected by WB after precipitation of proteins present in the extracellular medium of cortical cultures, as previously described *(27).* High-density DIV21 rat cortical neurons were incubated for 30 min in artificial cerebrospinal fluid (aCSF) composed of (in mM): 125 NaCl, 2.5 KCl, 2 CaCl_2_, 1.25 MgCl_2_, 11 glucose, 26.2 NaHCO_3_, 1 NaH_2_PO_4_, and 10 HEPES, pH 7.4). After incubation, the conditioned aCSF was collected and centrifuged at 500 x g or 7 min, to remove cell debris. The supernatant was collected and deoxycholic acid (Sigma Aldrich, USA: # D6750) was added to a final concentration of 0.015%. Tricholoroacetic acid (TCA) (Sigma Aldrich, USA: #T4885), pre-chilled to -20°C, was subsequently added to a final concentration of 20%. The samples were gently shaked and incubated for 30 min at -20°C, followed by centrifugation at 21000 x g for 30 min. The protein pellet was washed 3 times in ice-cold acetone (500 µL at -20°C) followed by centrifugation at 21000 x g for 10 min. The resulting pellet was briefly dried in a thermoblock at 60ºC for 2 min, taking care not to overdry. Finally, 40 µL of 2X Laemmli sample buffer was added.

**Immunoprecipitation (IP) of patient human IgGs to co-IP the endogenous ectodomain**

To assess whether CASPR2 IgGs could immunoprecipitate the soluble CASPR2 ectodomain, human IgGs (from patient or healthy controls) were added to DIV20 high density cortical neurons to a final concentration of 20 µg/mL. After 24h, the medium from DIV21 neurons was incubated with a mix of the protease inhibitors described above. To evaluate whether the autoantibodies are also bound to the soluble CASPR2 ectodomain in patients’ CSF, we used the same strategy and immunoprecipitated the soluble CASPR2 ectodomain from human CSF samples and controls that were diluted 4-fold to a final volume of 300 µL. Protein A Sepharose CL-4B (GE Healthcare, USA: #11514945) beads were used to capture the human IgG from the extracellular medium. The beads were prepared according to the manufacturer’s instructions, added to the collected media, and the suspension was left in orbital rotation ON at 4°C. On the subsequent day, the suspension was centrifuged at 2500 x g for 1 min at 4°C, and the first supernatant, corresponding to the nonbinding fraction, was collected. The pelleted beads were then resuspended and washed RIPA at 2500 x g for 1 min at 4°C for a total of 3 times. After the last wash, the human IgGs were released from the beads by denaturation in 2x concentrated Laemmli Sample Buffer and incubated at 95°C for 5 min. The beads suspension was centrifuged, and the supernatant was collected.

**Gel electrophoresis and Western Blot**

Protein samples or IP eluates were resolved by SDS-polyacrylamide gel electrophoresis (SDS-PAGE) in 9% polyacrylamide gels. Proteins were transferred onto a PVDF membrane (Millipore, USA: #IPVH00010) in Tris-Glycine buffer (TG; Tris 25 mM, Glycine 192 mM, pH 8.3) supplemented with 20% methanol by electroblotting at 300 mA ON at 4°C. The membranes were briefly washed in Tris-Buffered Saline (137 mM NaCl, 20 mM Tris-HCl, pH 7.6) containing 0.1% (v/v) Tween-20 (TBS-T) and then blocked in 5% (w/v) low-fat milk/TBS-T for 1h at RT. Membranes were probed with primary antibodies diluted in 5% (w/v) low-fat milk/TBS-T ON, at 4°C. On the subsequent day, membranes were washed with TBS-T for 30 min at RT and incubated with alkaline phosphatase-conjugated secondary antibodies for 1h at RT, followed by TBS-T washes. Membranes were then incubated with a chemifluorescent substrate (ECF, GE Heathcare, USA: #RPN5785) for 5 min at RT and scanned with the Chemidoc Imaging System (BioRad). Membrane signal was quantified using the ImageLab software (Bio-Rad, USA) under linear exposure conditions. If required, membranes were stripped with 0.2 M NaOH for 5 min, re-blocked and re-probed. Whenever necessary, total protein detection for normalization of protein loading was determined using the Revert 700 Total Protein Stain Kits for Western Blot Normalization (LICORBio, USA: #926-11016), per manufacturer instructions.

The presence of endogenous soluble CASPR2 ectodomain was probed with commercial antibodies directed against the N-terminal and C-terminal regions of CASPR2 and confirmed by a band of lower molecular weight compared to the whole-cell extracts and detected only with the N-terminal antibody. The absence of cell debris was confirmed by the lack of labelling for actin.

**Imaging in immunohistochemistry and immunocytochemistry experiments**

Immunohistochemistry (IHC) preparations of whole brain slices were imaged with a Plan Apochromat 2X/0.8 N.A. air objective in a Zeiss Axio Scan.Z1 microscope (Zeiss, Germany) or with a EC Plan-Neofluar 10X/0.3 air objective in a Widefield Zeiss Axio Imager Z2 microscope (Zeiss, Germany). ICC experiments used to validate virus titer were imaged with a lower amplification objective (Plan-Apochromat 40X/1.4 oil objective), in the same microscope as described above. All images were acquired with Zen Software.

**Live calcium imaging with GCaMP6f**

To assess the effects of human CASPR2 IgGs on calcium responses at the cell soma, neurons transduced with GCaMP6f AAVs were incubated with human IgGs (20 µg/ mL) at DIV14 or 15. After 24h, the neurons in coverslips were transferred to a warm insert with aCSF (in mM: 125 NaCl, 2.5 KCl, 2 CaCl_2_, 1.25 MgCl_2_, 11 Glucose, 26.2 NaHCO_3_, 1 NaH_2_PO_4_, and 10 HEPES, pH 7.4) and imaged using a Plan-Apochromat/0.8 objective in a Zeiss Cell Observer Spinning Disk at a time frame of 3 frames/second. After a basal recording of 60 seconds, cells were depolarized with 30 mM KCl to elicit a cytosolic Ca^2+^ signal, as previously published *(27).* For image analysis, regions of interest (ROIs) were drawn over the somas in the field of view using FIJI. To determine the peak amplitude of the calcium signal for each cell, the signal intensity of the ROI was corrected to the background signal and normalized to the average of the baseline recording. To determine the kinetics of the calcium signal decay, the data were further normalized to the peak amplitude and the plateau and fitted with an exponential function to obtain the t_1/2_ off in a systematic way using GraphPad Prism.

**Electrophysiology**

Whole-cell patch clamp recordings were used to assess the effect of patient IgGs on intrinsic excitability and synaptic transmission in DIV15-16 hDLX-mCherry^+^ (inhibitory) or hDLX-mCherry^-^ (excitatory) neurons incubated for 2h or 24h with the human IgGs. The coverslips were transferred to a recording chamber mounted on a fixed-stage inverted microscope (Zeiss Axio Observer.A1) and perfused with extracellular solution (140 mM NaCl, 2.4 mM KCl, 10 mM HEPES, 10 mM Glucose, 4 mM MgCl_2_, 4 mM CaCl_2_, pH 7.3, 300-310 mOsM) at a rate of 2-3 mL/min, at RT. Neurons were visualized under transmitted light using phase contrast, and inhibitory neurons were identified by mCherry fluorescence. Recordings were made using patch pipettes fabricated from borosilicate glass capillaries (Science Products GmbH, Germany) with a resistance of 3-5 MOhm when fill with an internal solution composed of (in mM):145 K-gluconate, 10 HEPES, 1 EGTA, 2 MgCl_2_, 2 MgATP, and 0.3 Na_3_GTP; pH 7.3, ~295 mOsM).

To determine membrane excitability, neurons were submitted to 500 ms current steps from -140 pA to + 480 pA increments, whereas the rheobase was estimated from a 500ms current ramp up to 500 pA (1pA/ms). The characteristics of single action potentials were determined from the first event triggered by a short square current pulse (5ms duration in 20pA increments). Spontaneous excitatory postsynaptic currents (sEPSCs) and spontaneous inhibitory postsynaptic currents (sIPSCs) were recorded under whole-cell voltage-clamp at -50 mV, allowing the simultaneous recording of excitatory (inward currents) and inhibitory events (outward currents) due to Cl^-^ influx in the absence of antagonists. Spontaneous PSCs were recorded for 5min in interspersed every minute with brief 10mV pulses to monitor access resistance during the recording. Signals were recorded with a HEKA EPC10 double USB patch clamp amplified, low-pass filtered at 2.9KHz, digitized at 20 KHz and acquired using PatchMaster Software (HEKA).

For all experiments, recordings were discarded if the holding current exceeded -200 pA. Recordings were also discarded if access resistance was higher than 25 MOhm or if it changed more than 20% over the course of the recording. Rheobase, AP count and kinetics were manually quantified with Igor Pro from Wavemetrics. The input resistance was obtained in voltage clamp by applying a 10 mV hyperpolarizing voltage step and calculating the steady-state resistance using Ohm’s law. The resting membrane potential (RMP) was determined as the membrane potential at zero current, estimated by averaging the voltage traces recorded during the 0 pA current step. Spontaneous Postsynaptic Currents (PSCs) events were quantified using a template matching algorithm (which was pre-trained with randomized recordings of these experiments) using Clampfit software (Axon Instruments). A total of 50 events were analyzed per recording.

**Microelectrode Array (MEA) assays in rat cortical neurons**

Spontaneous activity of cultured neurons incubated with either patient or healthy control IgGs was recorded using the Maestro Edge System (Axion Biosystems, USA). Prior to initiating a recording session, culture plates were visually inspected and wells with no cells, or with an uneven distribution of cells, were excluded *(50).* In addition, transduced cells were inspected for reporter protein expression to confirm viral infection. For the PMCA2 KD experiments, cells were incubated with either individual patient pIgGs or pooled patient samples. After acclimatizing the cultures for 10-15 min in the system in a controlled atmosphere of 95% air/5% CO_2_ at 37ºC, activity was recorded for 10 min before (baseline recording) and at several timepoints following incubation with the human IgGs. Data were sampled at 12.5 kHz and were digitized using the Axion Integrated Studio software (Axion Biosystems, GA, Atlanta).

MEAs recording data was quantified with user-defined algorithms using the Axis Navigator and Neuralmetrics tools for data extraction and analysis, respectively. Spike events were detected using an adaptive spike detection threshold of 6 SD for each electrode. Single-electrode bursts were defined as at least 5 consecutive spikes with interspike intervals shorter than 100 ms. Network bursts were defined using the interspike interval threshold algorithm considering at least 50 consecutive spikes across multiple electrodes with maximum interspike intervals of 10 ms and a minimum of 65% of participating electrodes. The results were expressed as a percentage of the baseline (prior to the incubation with IgGs). For the PMCA2 KD experiments, data were further normalized to Healthy Control pIgGs data.

**iNeuron MEA assays**

Neurons derived from hiPSCs (iNeurons) were generated from NPCs passaged no more than 6 times. NPCs were plated at 100,000 cells/cm² on Matrigel coated 6- well plates in Neural Progenitor Medium. After 24 hours the medium was switched to Forebrain Neuron Differentiation Medium. The medium, supplemented with 10 μM Y-27632, was fully replaced each day for 7 days. The resulting precursors were lifted using ACCUTASE™, spun down and then resuspended in DMEM/f12 before being counted. They were replated at 6 x 10⁴ cells per well onto PLO/laminin-coated 24-well MEA Plate-eco plates. Cells were then differentiated into neurons in STEMdiff™ Forebrain Neuron Maturation Medium. The medium was fully replaced every 3 to 4 days. Spontaneous activity was recorded for 10 minutes on a MED64 Presto system (Alpha Med Scientific, Osaka, Japan). Recordings were made weekly starting 2 weeks after differentiation began.

At week 7 after MEA plating, iNeurons were treated with healthy and CASPR2- patient sera at a 1:100 dilution in STEMdiff™ Forebrain Neuron Maturation Medium. Activity was recorded for 10 minutes at the start of each hour for a 6 hour period. A spike was counted when the signal passed ±5 standard deviations of the baseline noise. Baseline noise was measured during quiet periods. Bursts were defined as groups of spikes separated by intervals shorter than the mean inter-spike interval. Burst duration was set with the Poisson surprise method. Spikes and bursts were analyzed with MEA Symphony and Burst Analysis Table software (Alpha Med Scientific). An electrode was considered active when its mean firing rate was at least 5 spikes/min. Wells with fewer than 8 out of 16 active electrodes were excluded from analysis.

**Supplementary Tables**

**Table S1. Human CASPR2 pIgGs samples and IgG composition used in the MEA experiments with rat neurons**

| **ID** | **Age** | **Sex** | **Clinical symptoms** | **Cancer** | **Therapy response** | **Paired serum LCBA**  **end-point titer** | **CASPR2-IgG4/**  **Total** | **CASPR2-IgG4/**  **IgG1** |
| --- | --- | --- | --- | --- | --- | --- | --- | --- |
| Patient 1 | 72 | M | Personality change, amnesia, seizures, neuropathic pain | No | mRS 5 to 1 with corticosteroids and PLEX, later unexpected death | 1:6400 | unknown | unknown |
| Patient 2 | 73 | M | Peripheral nerve hyperexcitability, fatigue, no pain | No | mRS from 5 to 1 with azathioprine | 1:25600 | 0.25 | 3.45 |
| Patient 3 | 59 | F | Personality change, amnesia, seizures, neuropathic pain, peripheral nerve hyperexcitability, fatigue | Yes - Thymoma | mRS 4 to 2 with corticosteroids, IVIG and PLEX | 1:51200 | 0.26 | 27.61 |
| HC 1 | 73 | M | NA | NA | NA | Negative | NA | NA |
| HC 2 | 72 | M | NA | NA | NA | Negative | NA | NA |
| HC 3 | 40 | F | NA | NA | NA | Negative | NA | NA |

*Abbreviations: LCBA, live cell-based assay; F, female; HC, healthy control; IgG, immunoglobulin G; M, male; mRS, modified Rankin Scale; NA, not applicable; PLEX, plasma exchange, pIgGs,* Patient 1 plasma and pIgGs have been extensively characterized in prior studies.

**Table S2. Human sera samples from patients and control individuals used in the MEA experiments with human neurons**

| **ID** | **Age** | **Sex** | **Clinical symptoms** | **Cancer** | **Therapy response** | **Visual LCBA end point dilution** | **IgG1/**  **Total** | **IgG4/**  **Total** | **IgG1/4** | **IgG4/1** |
| --- | --- | --- | --- | --- | --- | --- | --- | --- | --- | --- |
| Patient 1 | 62 | M | Focal seizures, amnesia, behavioural, insomnia, fatigue | No | 2 to 0 mRS, Symptomatic therapy only | 25600 | NA | 0.27 | NA | NA |
| Patient 2 | 75 | M | Focal seizures, amnesia, behavioural, pain, dysautonomia, ataxia, fatigue | No | mRS from 4 to 2 with azathioprine | 12800 | 0.14 | 0.38 | 0.37 | 2.71 |
| HC 1 | 27 | M | NA | NA | NA | NA | NA | NA | NA | NA |

*Abbreviations: HC, Healthy Control; NA; Not Available or Not Applicable*

**Table S3. Human CSF samples from patients and control individuals**

| **ID** | **Sample Type** | **Visual LCBA end point dilution** | **IgG1/ Total** | **IgG4/ Total** | **IgG1/4** | **IgG4/1** |
| --- | --- | --- | --- | --- | --- | --- |
| Patient 1 | CASPR2 Patient CSF | 25600 | NA | 0.3306 | NA | NA |
| Patient 2 | CASPR2 Patient CSF | 12800 | 0.0190 | 0.2963 | 0.0583 | 15.6220 |
| Patient 3 | CASPR2 Patient CSF | 19200 | 0.0652 | 0.4547 | 0.1524 | 6.9761 |
| Patient 4 | CASPR2 Patient CSF | 25600 | 0.0090 | 0.3454 | 0.0260 | 38.3968 |
| Patient 5 | CASPR2 Patient CSF | 25600 | 0.0709 | 0.1840 | 0.4291 | 2.5962 |
| HC 1 | Control CSF (IIH) | 0 | NA | NA | NA | NA |
| HC 2 | Control CSF (IIH) | 0 | NA | NA | NA | NA |
| HC 3 | Control CSF (IIH) | 0 | NA | NA | NA | NA |

*Abbreviations: HC, Healthy Control; IIH, Idiopathic Intracranial Hypertension, NA; Not Available or Not Applicable*

**Table S4. Commercial antibodies**

| **Primary Antibodies** | **Application (Dilution)** | **Source** | **Reference** |
| --- | --- | --- | --- |
| Rabbit anti-GFP | ICC (1:500) | MBL | 598 |
| Chicken anti-MAP2 | ICC (1:5000) | Abcam | ab5392 |
| Rabbit anti-CNTNAP2 (C-terminal) | WB (1:1000) | Millipore | AB_5886 |
| Mouse anti-CNTNAP2 (N-terminal) | WB (1:250 or 1:500) | NeuroMab | 75-075 |
| Mouse anti-β-actin | WB (1:10000) | Sigma-Aldrich | A2228 |
| Rabbit anti-PMCA2 ATPase | WB (1:1000) | Thermo Fisher Scientific | PA1-915 |
| ***Secondary Antibodies*** |  |  |  |
| Goat anti-human IgG, Alexa Fluor® 568 | CBA ICC (1:500) | Life Technologies | A-21090 |
| Goat anti-human IgG, Alexa Fluor® 488 | CBA ICC (1:500) | Life Technologies | A-11013 |
| Donkey anti-rabbit IgG, Alexa Fluor® 488 | ICC (1:500) | Life Technologies | A-10042 |
| Goat anti-mouse IgG, Alexa Fluor® 568 | ICC (1:500) | Life Technologies | A-11004 |
| Goat anti-guinea pig IgG, Alexa Fluor® 647 | ICC (1:500) | Life Technologies | A-21450 |
| Goat anti-chicken IgG, Alexa Fluor® 647 | ICC (1:200) | Thermo Fisher Scientific | A-21449 |
| Goat anti-mouse IgG2a, Alexa Fluor® 488 | ICC (1:500) | Life Technologies | A-21241 |
| Goat anti-chicken IgG, AMCA | ICC (1:200) | Jackson ImmunoResearch | 103-155-155 |
| Goat anti-mouse IgG, alkaline phosphatase | WB (1:10000) | GE Healthcare | 115-055-003 |
| Goat anti-rabbit IgG, alkaline phosphatase | WB (1:10000) | GE Healthcare | 111-055-003 |
| Goat anti-human IgG, Biotinylated IgG | IHC (1:10000) | Vector Laboratories | BA-3000 |

*Abbreviations: ICC, immunocytochemistry; IHC, immunohistochemistry; WB, western blot*

**Table S5. DNA constructs**

| **DNA Constructs** | **Application** | **Source** |
| --- | --- | --- |
| *pEGFP-N1* | Live CBA | Clontech, France |
| *pCASPR2-EGFP* | Live CBA | Kind gift from Thomas Südhof, Standford University, USA |
| *pLV[Scramble shRNA]-mCherry-U6>rATP2B2* | LVs production | Kind gift from Peter Penzes, Northwestern University, USA |
| *pLV[PMCA2 shRNA]-mCherry-U6>rATP2B2* | LVs production | Kind gift from Peter Penzes, Northwestern University, USA |
| *pAAV.Syn.GCaMP6f.WPRE.SV40* | AAVs production | Addgene #153317 |
| *pAAV-hDLX-linker-mCherry* | AAVs production | Kind gift from Inna Slutsky, Tel Aviv University, Israel |
| *pAAV-mDlx-GFP-Fishell-1* | AAVs production | Addgene #83900 |
| *pDeltaF6* | AAVs production | Kind gift from Joris de Wit, VIB-KU Leuven Center for Brain & Disease Research, Belgium |
| *pRepCap* | AAVs production | Kind gift from Joris de Wit, VIB-KU Leuven Center for Brain & Disease Research, Belgium |
| *pPAX2* | LVs production | Kind gift from Joris de Wit, VIB-KU Leuven Center for Brain & Disease Research, Belgium |
| *pVSVG* | LVs production | Kind gift from Joris de Wit, VIB-KU Leuven Center for Brain & Disease Research, Belgium |

*Abbreviations: CBA, cell-based assay; LVs, lentiviruses; AAV, adeno-associated viruses.*

**Table S6. Experimental data and analyses for individual samples in MEA recordings**

| **Condition** | **N Exp.** | **N Wells** | **Statistical Test** |
| --- | --- | --- | --- |
| HC pIgGs | 7 | 37 | Two-way repeated measures ANOVA test |
| Patient 1 pIgGs | 7 | 40 | Two-way repeated measures ANOVA test |
| Patient 2 pIgGs | 7 | 36 | Two-way repeated measures ANOVA test |
| Patient 3 pIgGs | 7 | 40 | Two-way repeated measures ANOVA test |

*Abbreviations: HC, healthy control; pIgGs, purified Immunoglobulin G; N Exp., Number of Experiments.*

**Table S7. Experimental samples and analyses for spontaneous excitatory currents from inhibitory and excitatory neurons**

| **Exp.** | **Time** | **Cell Type** | **Condition** | **N Exp.** | **N Cells** | **Statistical Test** |
| --- | --- | --- | --- | --- | --- | --- |
| sEPSCs Amp | 2h | Excitatory | HC pIgGs | 6 | 23 | Mann-Whitney test |
|  |  |  | Patient pIgGs | 6 | 53 |  |
|  |  | Inhibitory | HC pIgGs | 6 | 20 |  |
|  |  |  | Patient pIgGs | 6 | 58 |  |
|  | 24h | Excitatory | HC pIgGs | 5 | 19 | Mann-Whitney test |
|  |  |  | Patient pIgGs | 5 | 43 |  |
|  |  | Inhibitory | HC pIgGs | 5 | 26 |  |
|  |  |  | Patient pIgGs | 5 | 61 |  |
| sEPSCs Freq | 2h | Excitatory | HC pIgGs | 6 | 23 | Mann-Whitney test |
|  |  |  | Patient pIgGs | 6 | 53 |  |
|  |  | Inhibitory | HC pIgGs | 6 | 20 |  |
|  |  |  | Patient pIgGs | 6 | 58 |  |
|  | 24h | Excitatory | HC pIgGs | 5 | 19 | Mann-Whitney test |
|  |  |  | Patient pIgGs | 5 | 43 |  |
|  |  | Inhibitory | HC pIgGs | 5 | 26 |  |
|  |  |  | Patient pIgGs | 5 | 61 |  |
| sEPSCs Amp x Freq | 2h | Excitatory | HC pIgGs | 6 | 23 | Mann-Whitney test |
|  |  |  | Patient pIgGs | 6 | 53 |  |
|  |  | Inhibitory | HC pIgGs | 6 | 20 |  |
|  |  |  | Patient pIgGs | 6 | 58 |  |
|  | 24h | Excitatory | HC pIgGs | 5 | 19 | Mann-Whitney test |
|  |  |  | Patient pIgGs | 5 | 43 |  |
|  |  | Inhibitory | HC pIgGs | 5 | 26 |  |
|  |  |  | Patient pIgGs | 5 | 61 |  |

*Abbreviations: sEPSCs, spontaneous excitatory postsynaptic currents; Freq, frequency; Amp, amplitude; HC, healthy control; N Exp., number of experiments.*

**Table S8. Experimental samples and analyses for spontaneous inhibitory currents from inhibitory and excitatory neurons**

| **Exp.** | **Time** | **Cell Type** | **Condition** | **N Exp.** | **N Cells** | **Statistical Test** |
| --- | --- | --- | --- | --- | --- | --- |
| sIPSCs Amp | 2h | Excitatory | HC pIgGs | 6 | 6 | Mann-Whitney test |
|  |  |  | Patient pIgGs | 6 | 24 |  |
|  |  | Inhibitory | HC pIgGs | 6 | 9 |  |
|  |  |  | Patient pIgGs | 6 | 22 |  |
|  | 24h | Excitatory | HC pIgGs | 5 | 7 | Mann-Whitney test |
|  |  |  | Patient pIgGs | 5 | 25 |  |
|  |  | Inhibitory | HC pIgGs | 5 | 11 |  |
|  |  |  | Patient pIgGs | 5 | 39 |  |
| sIPSCs Freq | 2h | Excitatory | HC pIgGs | 6 | 6 | Mann-Whitney test |
|  |  |  | Patient pIgGs | 6 | 24 |  |
|  |  | Inhibitory | HC pIgGs | 6 | 9 |  |
|  |  |  | Patient pIgGs | 6 | 22 |  |
|  | 24h | Excitatory | HC pIgGs | 5 | 7 | Mann-Whitney test |
|  |  |  | Patient pIgGs | 5 | 25 |  |
|  |  | Inhibitory | HC pIgGs | 5 | 11 |  |
|  |  |  | Patient pIgGs | 5 | 39 |  |
| sIPSCs Amp x Freq | 2h | Excitatory | HC pIgGs | 6 | 6 | Mann-Whitney test |
|  |  |  | Patient pIgGs | 6 | 24 |  |
|  |  | Inhibitory | HC pIgGs | 6 | 9 |  |
|  |  |  | Patient pIgGs | 6 | 22 |  |
|  | 24h | Excitatory | HC pIgGs | 5 | 7 | Mann-Whitney test |
|  |  |  | Patient pIgGs | 5 | 25 |  |
|  |  | Inhibitory | HC pIgGs | 5 | 11 |  |
|  |  |  | Patient pIgGs | 5 | 39 |  |

*Abbreviations: sIPSCs, spontaneous inhibitory postsynaptic currents; Freq, frequency; Amp, amplitude; HC, healthy control; N Exp., number of experiments*

**Table S9. Experimental samples and analyses of passive membrane properties from excitatory and inhibitory neurons**

| **Exp.** | **Time** | **Cell Type** | **Condition** | **N Exp.** | **N Cells** | **Statistical Test** |
| --- | --- | --- | --- | --- | --- | --- |
| Capacitance | 2h | Excitatory | HC pIgGs | 6 | 25 | Mann-Whitney test |
|  |  |  | Patient pIgGs | 6 | 63 |  |
|  |  | Inhibitory | HC pIgGs | 6 | 26 |  |
|  |  |  | Patient pIgGs | 6 | 67 |  |
|  | 24h | Excitatory | HC pIgGs | 6 | 27 | Mann-Whitney test |
|  |  |  | Patient pIgGs | 6 | 49 |  |
|  |  | Inhibitory | HC pIgGs | 6 | 32 |  |
|  |  |  | Patient pIgGs | 6 | 63 |  |
| Input Resistance | 2h | Excitatory | HC pIgGs | 6 | 25 | Mann-Whitney test |
|  |  |  | Patient pIgGs | 6 | 61 |  |
|  |  | Inhibitory | HC pIgGs | 6 | 27 |  |
|  |  |  | Patient pIgGs | 6 | 66 |  |
|  | 24h | Excitatory | HC pIgGs | 6 | 25 | Mann-Whitney test |
|  |  |  | Patient pIgGs | 6 | 51 |  |
|  |  | Inhibitory | HC pIgGs | 6 | 32 |  |
|  |  |  | Patient pIgGs | 6 | 66 |  |
| RMP | 2h | Excitatory | HC pIgGs | 6 | 23 | Mann-Whitney test |
|  |  |  | Patient pIgGs | 6 | 58 |  |
|  |  | Inhibitory | HC pIgGs | 6 | 25 |  |
|  |  |  | Patient pIgGs | 6 | 62 |  |
|  | 24h | Excitatory | HC pIgGs | 6 | 27 | Unpaired t-test |
|  |  |  | Patient pIgGs | 6 | 61 |  |
|  |  | Inhibitory | HC pIgGs | 6 | 24 |  |
|  |  |  | Patient pIgGs | 6 | 49 |  |

*Abbreviations: RMP, resting membrane potential; HC, healthy control; N Exp., number of experiments.*

**Table S10. Experimental samples and analyses for action potential characteristics from excitatory and inhibitory neurons**

| **Exp.** | **Time** | **Cell Type** | **Condition** | **N Exp.** | **N Cells** | **Statistical Test** |
| --- | --- | --- | --- | --- | --- | --- |
| Threshold | 2h | Excitatory | HC pIgGs | 6 | 19 | Mann-Whitney test |
|  |  |  | Patient pIgGs | 6 | 41 |  |
|  |  | Inhibitory | HC pIgGs | 6 | 17 |  |
|  |  |  | Patient pIgGs | 6 | 47 |  |
|  | 24h | Excitatory | HC pIgGs | 6 | 19 | Mann-Whitney test |
|  |  |  | Patient pIgGs | 6 | 41 |  |
|  |  | Inhibitory | HC pIgGs | 6 | 29 |  |
|  |  |  | Patient pIgGs | 6 | 59 |  |
| Rise Time | 2h | Excitatory | HC pIgGs | 6 | 19 | Mann-Whitney test |
|  |  |  | Patient pIgGs | 6 | 41 |  |
|  |  | Inhibitory | HC pIgGs | 6 | 17 |  |
|  |  |  | Patient pIgGs | 6 | 47 |  |
|  | 24h | Excitatory | HC pIgGs | 6 | 23 | Mann-Whitney test |
|  |  |  | Patient pIgGs | 6 | 44 |  |
|  |  | Inhibitory | HC pIgGs | 6 | 29 |  |
|  |  |  | Patient pIgGs | 6 | 59 |  |
| Half-width time | 2h | Excitatory | HC pIgGs | 6 | 19 | Mann-Whitney test |
|  |  |  | Patient pIgGs | 6 | 41 |  |
|  |  | Inhibitory | HC pIgGs | 6 | 17 |  |
|  |  |  | Patient pIgGs | 6 | 47 |  |
|  | 24h | Excitatory | HC pIgGs | 6 | 23 | Mann-Whitney test (24h) and Unpaired t-test (2h) |
|  |  |  | Patient pIgGs | 6 | 44 |  |
|  |  | Inhibitory | HC pIgGs | 6 | 29 |  |
|  |  |  | Patient pIgGs | 6 | 59 |  |
| Decay time | 2h | Excitatory | HC pIgGs | 6 | 19 | Mann-Whitney test |
|  |  |  | Patient pIgGs | 6 | 41 |  |
|  |  | Inhibitory | HC pIgGs | 6 | 17 |  |
|  |  |  | Patient pIgGs | 6 | 47 |  |
|  | 24h | Excitatory | HC pIgGs | 6 | 23 |  |
|  |  |  | Patient pIgGs | 6 | 43 |  |
|  |  | Inhibitory | HC pIgGs | 6 | 29 |  |
|  |  |  | Patient pIgGs | 6 | 59 |  |
| AP Peak amplitude | 2h | Excitatory | HC pIgGs | 6 | 19 | Mann-Whitney test |
|  |  |  | Patient pIgGs | 6 | 41 |  |
|  |  | Inhibitory | HC pIgGs | 6 | 17 |  |
|  |  |  | Patient pIgGs | 6 | 47 |  |
|  | 24h | Excitatory | HC pIgGs | 6 | 23 |  |
|  |  |  | Patient pIgGs | 6 | 44 |  |
|  |  | Inhibitory | HC pIgGs | 6 | 29 |  |
|  |  |  | Patient pIgGs | 6 | 59 |  |
| AP amplitude | 2h | Excitatory | HC pIgGs | 6 | 23 | Mann-Whitney test |
|  |  |  | Patient pIgGs | 6 | 44 |  |
|  |  | Inhibitory | HC pIgGs | 6 | 17 |  |
|  |  |  | Patient pIgGs | 6 | 47 |  |
|  | 24h | Excitatory | HC pIgGs | 6 | 23 |  |
|  |  |  | Patient pIgGs | 6 | 44 |  |
|  |  | Inhibitory | HC pIgGs | 6 | 29 |  |
|  |  |  | Patient pIgGs | 6 | 59 |  |
| AHP | 2h | Excitatory | HC pIgGs | 6 | 12 | Mann-Whitney test |
|  |  |  | Patient pIgGs | 6 | 28 |  |
|  |  | Inhibitory | HC pIgGs | 6 | 17 |  |
|  |  |  | Patient pIgGs | 6 | 40 |  |
|  | 24h | Excitatory | HC pIgGs | 6 | 14 |  |
|  |  |  | Patient pIgGs | 6 | 31 |  |
|  |  | Inhibitory | HC pIgGs | 6 | 29 |  |
|  |  |  | Patient pIgGs | 6 | 57 |  |

*Abbreviations: AP, action potential; AHP, afterhyperpolariztion; HC, healthy control; N Exp., number of experiments.*
